# small RNA can move long distances through plant vasculature to influence gene expression in shoot apical meristems

**DOI:** 10.1101/2021.09.20.460861

**Authors:** Mark A. A. Minow, Viktoriya Coneva, Victoria Lesy, Max Misyura, Joseph Colasanti

## Abstract

In plants, small RNA (sRNA) can regulate gene expression via post transcriptional gene silencing (PTGS) or through RNA-directed DNA methylation (RdDM) leading to transcriptional gene silencing (TGS). sRNA is mobile throughout the plant, with movement occurring short distances from cell-to-cell as well as long distances through the vasculature via phloem trafficking. The range of long-distance sRNA mediated signaling from the vasculature to the shoot apical meristem (SAM) is not clear. To investigate this, two independent transgenic approaches were used to examine trafficking of phloem-expressed sRNA to the SAM in *Arabidopsis thaliana*. First, the phloem companion-cell specific promoter *SUC2* was used to drive expression of an inverted repeat complementary to *FLOWERING LOCUS D* (*FD)*, a flowering time regulator expressed exclusively in the SAM. In a separate experiment, the *SUC2* promoter was used to express an artificial microRNA (aMiR) designed to target a synthetic *CLAVATA3* (*CLV3*) target in the SAM stem cells. Both systems provide evidence of a phloem-to-SAM sRNA communication axis connecting distal regions of the plant to the stem cells of the SAM, which ultimately gives rise to all shoot tissues, including gametes. Thus, phloem-to-SAM sRNA movement defines an important link between sRNA expressed in distal regions of the plant and the growing shoot. Importantly, phloem-to-SAM sRNA trafficking may allow somatic sRNA to direct SAM RdDM, fixing transient sRNA expression events into stable epigenetic changes.

## Introduction

Plant shoots grow from a small collection of stem cells located within the shoot apical meristem (SAM). This niche of stem cells is established during embryogenesis and maintained during somatic growth to provide progenitor cells that differentiate into all above-ground tissues, including the flowers that generate male and female gametophytes. In *Arabidopsis thaliana* the stems cell population in the SAM is partly controlled by a feedback loop involving *CLAVATA3* (*CLV3*) (Clark *et al*. 1995; Schoof *et al*. 2000). Stems cells in the central zone express *CLV3*, which encodes a non-autonomous signal peptide that moves to the subtending cells to inhibit meristem proliferation (Fletcher 1999; Rojo *et al*. 2002). Loss of *CLV3* activity results in more cells produced in the SAM, giving rise to larger organs and additional floral organs (Clark *et al*. 1995). *CLV3* expression is balanced by an endogenous compensation loop (Schoof *et al*. 2000; Rodriguez-Leal *et al*. 2019). In *clv3* mutants that produce non-functional transcript, this compensation results in a dramatic upregulation of *CLV3* expression (Rodriguez-Leal *et al*. 2019). Due to the CLV3 signaling compensation loop, a strong reduction in *CLV3* expression (below 33% of Wt levels) is needed to manifest a ‘clv’ mutant phenotype (Müller *et al*. 2006). For example, only 4/35 inducible *35S* RNAi lines produced sufficient *CLV3* knockdown to elicit ‘clv’ phenotypes (Reddy and Meyerowitz 2005). The CLV3 regulatory circuit is conserved in higher plants, and subtle variations in activity alter the size and shape of the shoot organs, contributing to the extensive diversity of higher plants, including crops (Somssich *et al*. 2016).

Key gene regulation changes at the SAM herald shoot developmental transitions. One critical developmental transition at the SAM is the switch from vegetative to reproductive growth, which affects fitness in natural environments and agronomic yield. The initiation of SAM reproductive development is influenced by the environment; for example, long-day (LD) photoperiods hasten flowering time in Arabidopsis is hastened (Andrés and Coupland 2012). *FD* is an important regulator of Arabidopsis flowering that is constitutively expressed in the SAM, and *fd* mutants exhibit delayed flowering under LDs (Abe *et al*. 2005; Wigge *et al*. 2005). To trigger flowering, the FD transcription factor interacts with mobile floral inductive signals that are produced in leaf companion cells and transmitted to the SAM via the phloem. Leaves perceive long-day photoperiods and subsequently express *FLOWERING LOCUS T* (*FT*), a phloem mobile signal (florigen) that promotes flowering through interactions with FD (Abe et al. 2005; Wigge et al. 2005; Corbesier et al. 2007). The phloem also traffics other information molecules throughout the plant to integrate development with the environment (Turnbull and Lopez-Cobollo 2013; Kehr and Kragler 2018). In addition to protein factors, the phloem transports small RNA (sRNA) that can act as important gene regulators (Buhtz *et al*. 2008; Pant *et al*. 2008).

sRNAs are known to mediate gene silencing in diverse eukaryotes. Plant sRNAs are the products of DICER-LIKE (DCL) enzymatic processing of diverse double stranded RNA (dsRNA) substrates, producing ∼20-24 nucleotide (nt) sRNA duplexes (Borges and Martienssen 2015). *HUA ENHANCER 1* (*HEN1*) then methylates the 3’ end of sRNA to promote RNA stability with *hen1* mutants under-expressing sRNA of all sizes (Yang *et al*. 2006). Mature sRNAs interact with ARGONAUTE (AGO) proteins to repress mRNA activity through post-transcriptional gene silencing (PTGS) or by repressing transcription through transcriptional gene silencing (TGS) (Vaucheret 2008). PTGS typically involves 21/22nt sRNA that target complementary mRNA molecules for cleavage or translational inhibition (Baumberger and Baulcombe 2005; Brodersen *et al*. 2008; Reis *et al*. 2015). TGS at target genes has been associated with transgenerationally heritable DNA methylation changes that are triggered by the RNA directed DNA methylation (RdDM) cycle (Matzke and Mosher 2014). Although not fully understood, non-canonical RdDM links PTGS with TGS; that is, silencing can progress from repressing translation to preventing transcription (Cuerda-Gil and Slotkin 2016).

MicroRNAs (miRNAs) are typically 21nt in length and serve as evolutionarily conserved master regulators of many processes, including leaf development, aging, the floral transition and environmental sensing (Fujii *et al*. 2005; Chitwood *et al*. 2009; Wu *et al*. 2009; Koyama *et al*. 2017). miRNAs are produced from transcripts encoding imperfectly complementary hairpin loops and typically elicit PTGS (Borges and Martienssen 2015). However, miRNAs can alternatively trigger TGS either directly or by eliciting secondary siRNA biogenesis (transitivity), often through RNA-DEPENDENT RNA POLYMERASE6 (RDR6), which then initiate RdDM (Chellappan et al. 2010; Wu et al. 2010; Creasey et al. 2014). Small interfering RNA (siRNA) typically originate from fully complementary dsRNA like large inverted repeats (Borges and Martienssen 2015). In contrast to miRNA, siRNA embody a collection of sRNA of diverse sizes and regulatory actions.

Some plant sRNAs have been shown to regulate genes non-autonomously by moving short distances from cell to cell through plasmodesmata (Voinnet et al. 1998; Dunoyer et al. 2007; Smith et al. 2007; Melnyk et al. 2011b; Vatén et al. 2011; Rosas-Diaz et al. 2018). This local diffusion facilitates the morphogen-like behavior of several miRNAs by creating developmental gradients (Chitwood *et al*. 2009). Mobile sRNA can also elicit transitivity in recipient cells (Himber et al. 2003; Allen et al. 2005; Yoshikawa et al. 2005; Donaire et al. 2008; Liang et al. 2012). These secondary siRNAs move also, initiate transitivity, and silence targets at a short distance from the initial signal. In this way, secondary siRNAs can establish organism-wide gene silencing by cycling between local sRNA movement and sRNA signal amplification (Palauqui et al. 1997; Voinnet and Baulcombe 1997; Liang et al. 2012). Additionally, plant sRNA can move long distances via phloem sieve tubes without requiring continual RDR amplification (Molnar *et al*. 2010; Melnyk *et al*. 2011a). Phloem-localized sRNA is composed of all size classes, including siRNA and miRNAs that respond to environmental conditions (Buhtz *et al*. 2008; Pant *et al*. 2008). Long-distance phloem movement occurs across graft junctions and can move into recipient tissues such as root meristems and fully vascularized flowers (Melnyk et al. 2011a; Zhang et al. 2014; Li et al. 2021). However, long distance movement of phloem-derived sRNA into the SAM remains ambiguous (Voinnet *et al*. 1998; Brosnan and Voinnet 2011).

To study the potential movement of long-distance sRNA signals, parallel transgenic approaches were used here to determine whether Arabidopsis sRNA can move phloem-to-SAM to regulate gene expression. In one transgenic system, phloem expression of an inverted repeat targeted the *FD* flowering time regulatory gene that is localized throughout the SAM. This analysis showed that sRNA derived from a phloem expressed inverted repeat can move to the SAM and inhibit *FD* activity, resulting in delayed flowering. In a second system aimed at targeting the small population of stem cells in the SAM, a phloem-expressed artificial miRNA (aMiR) was able to repress a synthetic *CLV3* target in the SAM. This analysis provided additional evidence that sRNA can move phloem-to-SAM and, specifically, that mobile sRNA can enter an isolated subdomain with the SAM central zone. This phloem-to-SAM sRNA signaling suggests that somatic sRNA could influence developmental decisions originating in the SAM. Moreover, since the SAM gives rise to future gametophytes, phloem-to-SAM sRNA signaling might trigger epigenetic changes allowing somatic sRNA to have transgenerational consequences.

## Results

### Phloem-expressed sRNA can act as an ‘anti-florigen’

To determine whether phloem-derived sRNA can influence gene regulation in the SAM we utilized two components: a phloem-expressed sRNA signal and a SAM specific target. In Arabidopsis, the *SUCROSE SYMPORTER 2* (*pSUC2*) promoter is a well characterized promoter that drives specific expression in companion cells, and has been used in previous Arabidopsis phloem-to-SAM movement studies (Truernit and Sauer 1995; Corbesier et al. 2007). We independently verified the phloem specificity of this promoter by expressing a *pSUC2::β-GLUCURONIDASE:GREEN FLUORESCENT PROTEIN* (*GUS:GFP*) cassette, which exhibited phloem specific reporter activity subtending the SAM in all independent lines (n=3; Figure 1; Figure S1). *FD* was selected as the endogenous target, as it is constitutively expressed in the SAM, and loss of *FD* function causes delayed flowering that is observed easily under LD growth conditions (Abe et al. 2005; Wigge et al. 2005). To test phloem-to-SAM sRNA movement, *pSUC2* was used to drive expression of an inverted repeat homologous to *FD* (hereafter *pSUC2::FDi*) (Figure 1a). As a control to ensure the *FDi* repeat could knockdown *FD* without movement, the *FDi* repeat was cloned under the control of the native *FD* promoter (*pFD::FDi*). We hypothesized that, if sRNA produced by the *pSUC2::FDi* construct can reach the SAM and inhibit *FD* expression, *pSUC2::FDi* transgenics should flower later under LDs.

**Figure 1.**
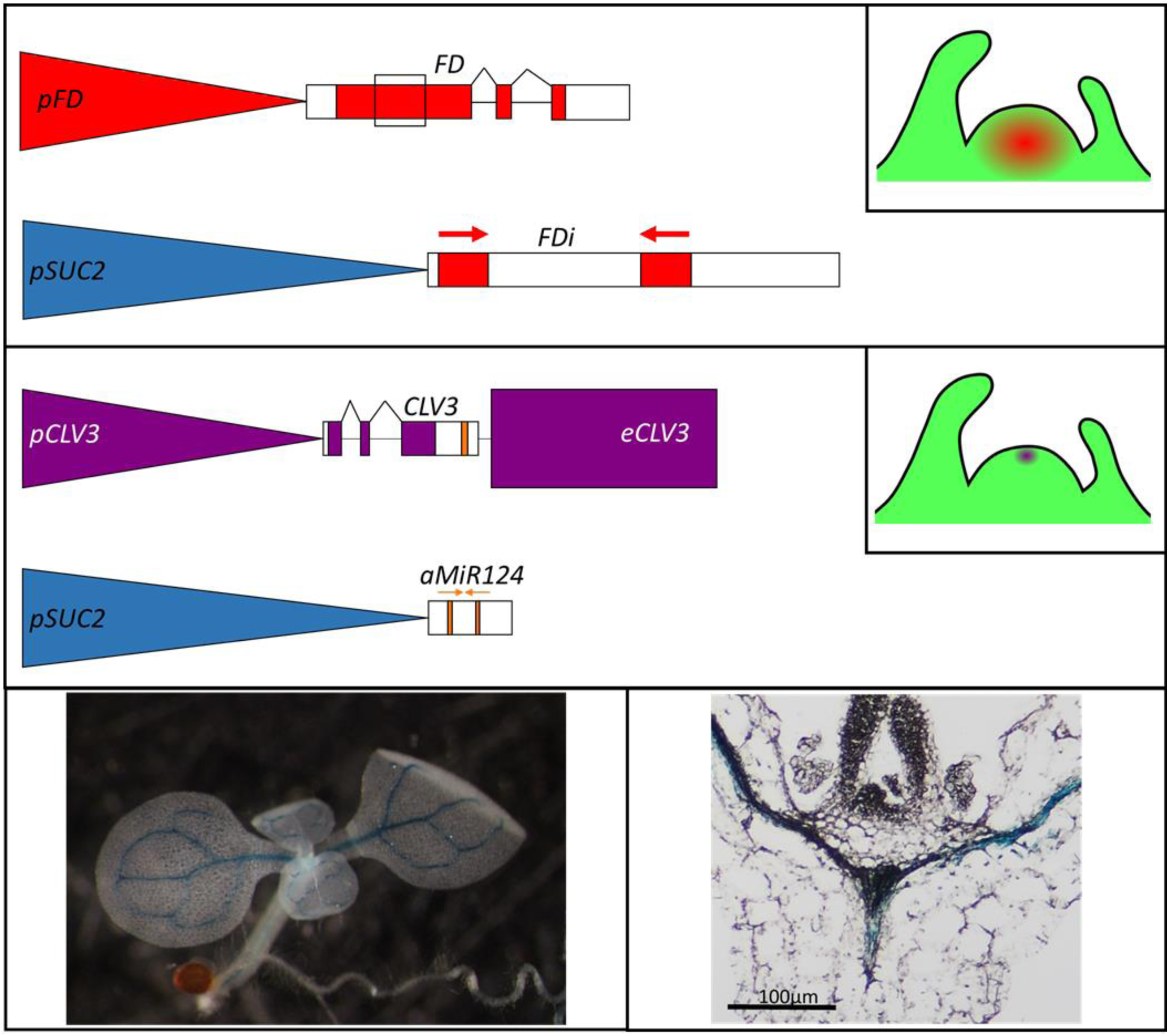
The two Arabidopsis transgenic systems devised to test phloem-to-SAM sRNA silencing. (**A**) The first system uses a transgenic sRNA source and an endogenous SAM target. The transgene has the *SUC2* promoter (*pSUC2*; blue) driving phloem expression of an inverted repeat (*FDi*) homologous to *FD* (red). If the phloem expressed *pSUC2::FDi* can repress *FD* in the SAM, then the floral transition should be delayed under long day photoperiods. Top Inset: Depiction of where *FD* is expressed in the SAM. The region in the first *FD* exon homologous to the *FDi* cassette is denoted by a black box. (**B**) The second system uses a transgenic sRNA source and SAM target. A site complementary to *MmuMiR124* (gold) was inserted into the 3’ UTR of the *CLV3* (purple) transcript. This modified *CLV3* transcript was cloned with the native *CLV3* promoter (*pCLV3*) and 3’ enhancer (*eCLV3*). This *CLV3* transgene (*pCLV3::CLV3: MmuMiR124’::eCLV3*; abbreviated to *C*) was then used to rescue *clv3-2* mutants. After selecting for stable homozygous *C* lines, a second transgene was stacked into these plants. This transgene used *pSUC2* to expresses an *aMiR* consisting of the mature *MmuMiR124* sequence in the Arabidopsis *MiR319a* precursor transcript. If the *pSUC2::aMmuMiR124* transgene (abbreviated to *S*) can silence *C* expression, then the stacked transgenics should show enlarged SAMs and an increase in floral organ numbers. **Middle Inset**: Depiction of where *CLV3* is expressed in the SAM. (**C&D**) A *pSUC2::GUS:GFP* transcriptional reporter was used to confirm *pSUC2* vein specificity in Col. Both cleared whole mounts (**C**) and paraffin wax embedded and sectioned tissues (**D**) exhibited GUS reporter staining constricted to the veins, distal to the shoot apex. The black scale bar denotes the length corresponding to 500 base pairs (bp) of DNA.

Twelve and 17 independent *pSUC2::FDi* lines were created in Landsberg *erecta* (Ler) and Columbia 0 (Col) accessions, respectively. Additionally, 5 and 17 independent lines of the *pFD::FDi* control were created in Ler and Col, respectively. All T2 lines screened displayed variable levels of delayed flowering (Figure S2). Three Col *pSUC2::FDi* plants with a pronounced delay were selected for further analysis. In progressive generations (up to T7), individuals of these three lines exhibited delayed flowering under LDs that was inherited with the transgene as a dominant trait (Figure 2). The delayed flowering in *pSUC2::FDi* lines was not as strong as that produced by *pFD::FDi* lines nor by the *fd-5* mutants (Figure 2). *pSUC2::FDi*, *pFD::FDi* or *fd-5* plants grown under non-inductive short days flowered at the same time as Col Wt (Figure S3). Quantitative real-time PCR (qPCR) showed that the severity of the floral delay corresponded to *FD* mRNA levels; *pSUC2::FDi* lines exhibited reduced *FD* expression, but the *pFD::FDi* lines and the *fd-5* mutant both had lower *FD* expression (Figure 3). Furthermore, crossing the *pSUC2::FDi* transgene into *fd-5* mutant plants revealed no additive delay of flowering, indicating the knockdown was specific to *FD* activity (Figure S4). These data indicate that the *pSUC2::FDi* transgene does repress *FD* over a distance and that this distal repression is less effective than proximal RNAi or genetic disruption of *FD*.

**Figure 2.**
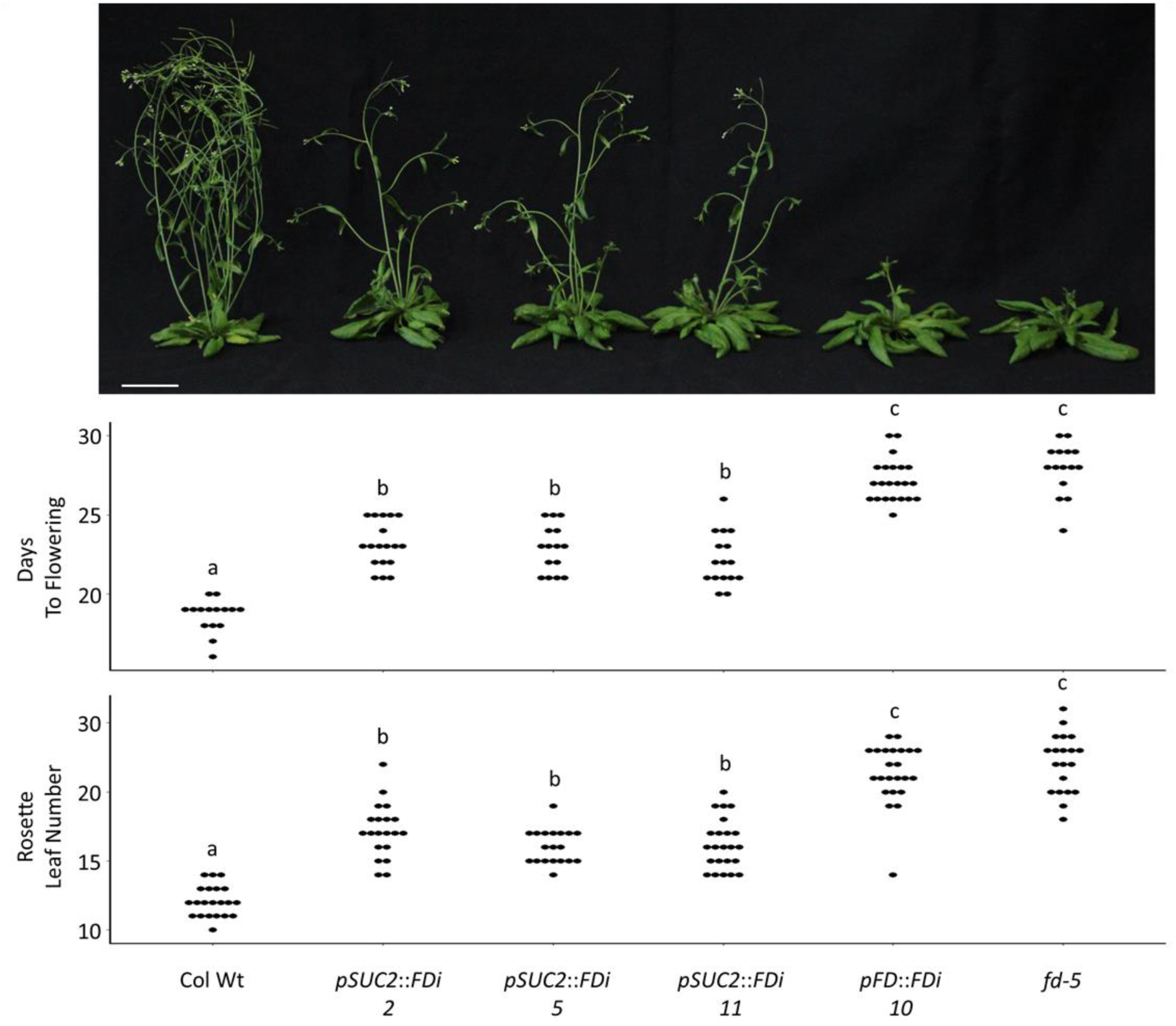
The *pSUC2::FDi* transgene delays the floral transition under long day photoperiods. Plants containing the *pSUC2::FDi* transgene produce more rosette leaves and flower significantly later than the Wt control. Three independent *pSUC2::FDi* lines are shown. The *pSUC2::FDi* floral delay was not as strong as that observed by expressing the same *FDi* cassette under the *FD* promoter nor the *fd-5* mutant. Each dot represents one individual for each respective genotype.

**Figure 3.**
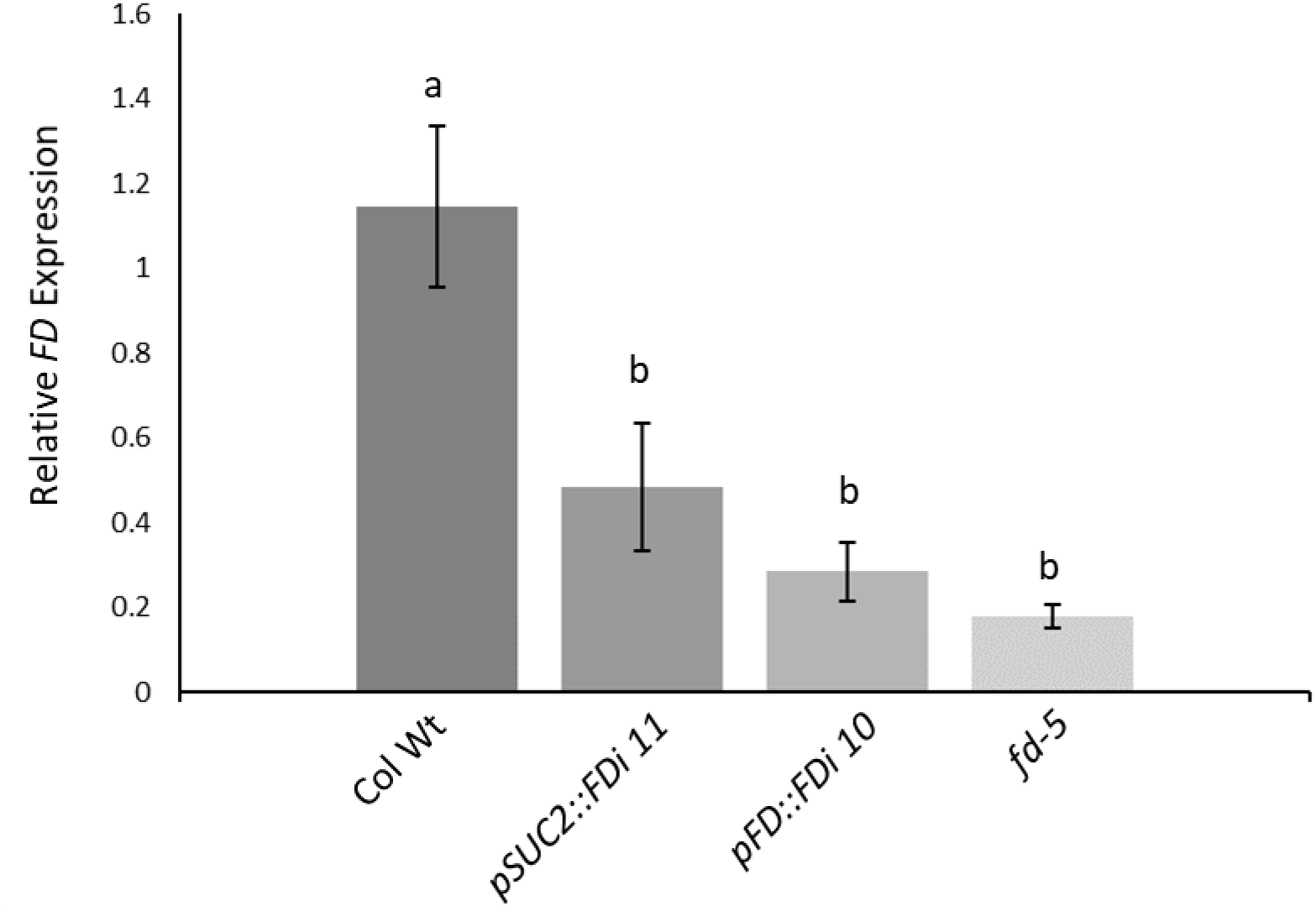
qPCR of apically enriched tissues indicate the *pSUC2::FDi* transgene reduces *FD* expression. The measurement of FD expression from a representative line indicated the *pSUC2::FDi* transgene reduced FD expression to ∼50% that of WT levels. Although not statistically significant, *FD* expression in the representative *pFD::FDi* line and *fd-5* mutants trended towards lower expression than *pSUC2::FDi 11*. *FD* expression is normalized to *KNAT1*. Letters denote statistically similar (p>0.05) groups, as determined by ANOVA with *post hoc* Tukey’s HSD test. Error bars indicate +/- standard error. Scale bar denotes 5 cm.

### Defects in sRNA production abrogate the *pSUC2::FDi* effect

To provide evidence that the *pSUC2::FDi* floral delay was sRNA driven, the line carrying this construct with the most consistent floral delay was crossed into three lines that were deficient in genes required for sRNA biogenesis: *HEN1, RDR6* and *HASTY (HST)*. If the activity of these sRNA related genes is required for *pSUC2::FDi* function, the mutant backgrounds should abrogate the floral delay caused by the *pSUC2::FDi* transgene. *pSUC2::FDi hen1-6* plants showed no delay in flowering compared to *hen1-6* alone, indicating sRNA facilitates knockdown of *FD* (Figure S5). Similarly, *pSUC2::FDi rdr6-15* plants displayed equivalent flowering time to *rdr6-15* mutants (Figure S5). The *rdr6-15* suppression of the *pSUC2::FDi* floral delay suggests the involvement of RDR6 amplification in phloem-to-SAM sRNA silencing. The *pSUC2::FDi* gene was also crossed with two different alleles of *HASTY*, an importin/exportin homolog that is reported to be involved in nuclear sRNA export (Peragine et al. 2004; Park et al. 2005). Again, the combination of *pSUC2::FDi* with either the *hst-6* or *hst-15* allele suppressed the late flowering phenotype (Figure S5). However, the loss of *HST* function inhibits the actions of floral repressor *miR156,* thus causing an acceleration of the floral transition (Wu and Poethig 2006). Therefore, it is unclear whether the lack of floral delay in the *pSUC2::FDi hst^-^* plants is due to the failure of *FDi* sRNA act at the SAM or because of aberrant *hst^-^ miR156* levels. Taken together, these data support phloem-to-SAM mobile sRNA eliciting delayed flowering in *pSUC2::FDi* plants.

### Phloem-expressed artificial miRNA influences gene expression in SAM stem cells

To investigate more precisely the movement of sRNA into the SAM, we constructed a second transgenic system to determine whether phloem-derived sRNA can move into the SAM and affect the activity of *CLV3*, a gene expressed specifically in the small population of stem cells in the apical region of the meristem. *CLV3* was chosen as a target for sRNA knockdown as it is expressed exclusively in stem cells (Schoof et al. 2000; Brand et al. 2002). To test this idea, the native *CLV3* gene was cloned with its native promoter (*pCLV3*) and the 3’ enhancer element (*eCLV3*) that is required for central zone expression (Brand *et al*. 2002). The 3’ UTR of this *CLV3* transgene was modified to contain a site complementary to mouse microRNA *MmuMiR124. MmuMiR124* was chosen as it has no plant homolog and no complementary sequences were detected in the Arabidopsis genome (Ghosh Dastidar *et al*. 2016; Wójcik *et al*. 2018). This recombinant construct, *pCLV3::CLV3:MmuMiR124’::eCLV3*, is hereafter abbreviated to ‘*C’* (Figure 1b). The sRNA generating component of this system is an *aMiR* consisting of the Arabidopsis *MiR319a* precursor transcript with the active miRNA site replaced with the *MmuMiR124* sequence expressed by *pSUC2*. This construct, *pSUC2::aMmuMiR124*, is hereafter abbreviated to ‘*S’* (Figure 1b). We hypothesize that, if sRNA produced from *S* in the phloem can reach the mRNA expressed from *C* in the stem cells, it should repress *C* expression (*S->C* silencing) and, consequently, produce a similar phenotype to that exhibited by *clv3-2* mutants. In addition to the previously reported increase in flower and floral organs, we observed that *clv3-2* plants produce additional rosette leaves (Figure 4C; Figure S6).

**Figure 4.**
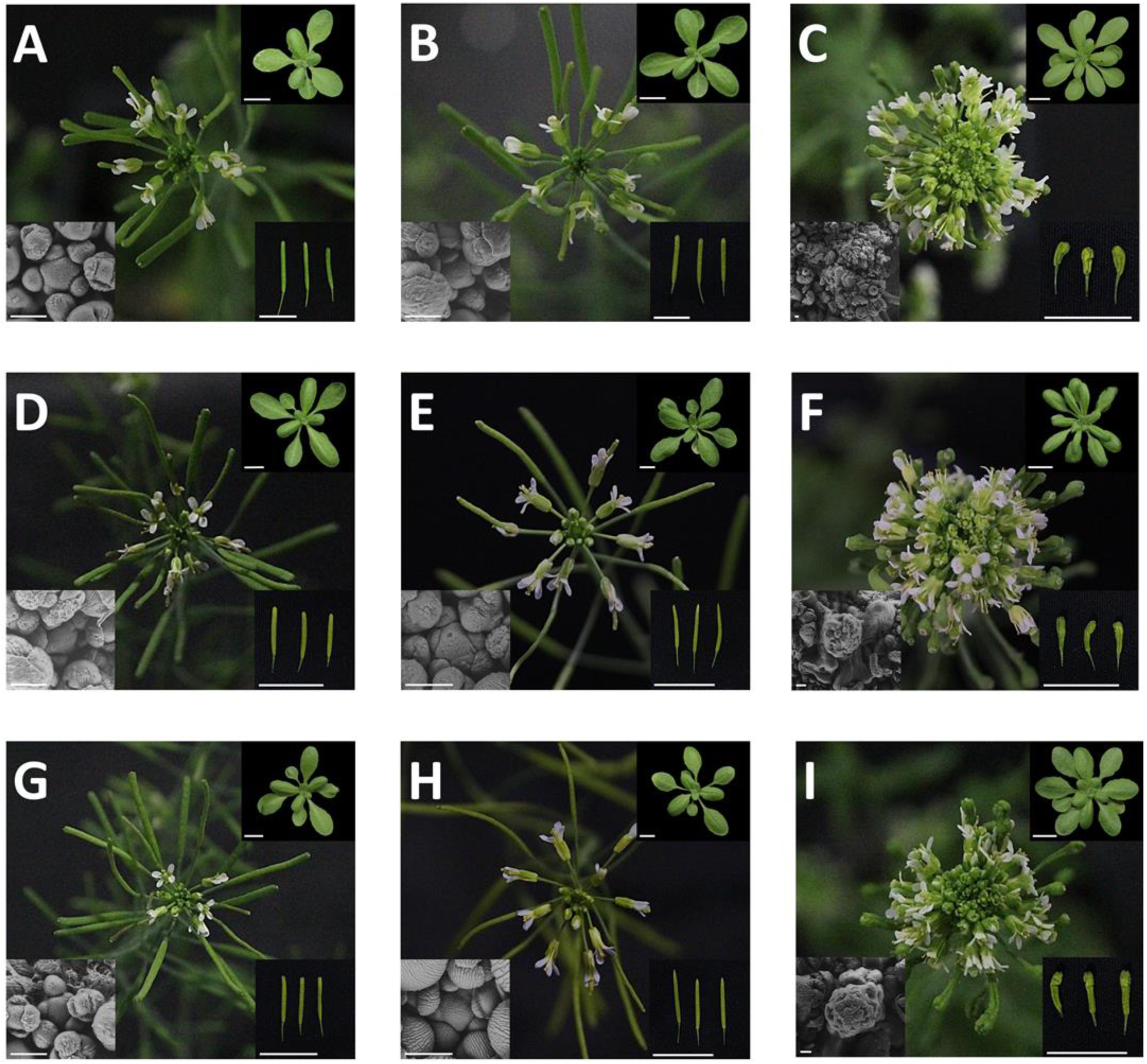
*S->C* transgenics shows variable silencing with some strong ‘clv’ events. Representative images representing the inflorescence (center), rosette leaves (upper right; scale = 1cm), silique (lower right; scale = 1cm) and SAM (bottom left scale = 100μm) phenotypes for the following groups: (A) Ler Wt (B) Ler *pSUC2::aMiR124* (C) *clv3-2* (D) *C9* (E) *S->C 9-3* ‘Wt’ (F) *S->C 9-3* ‘clv’, (G) *C10*, (H) *S->C 10-5* ‘Wt’, (I) *S->C 10-5* ‘clv.’ Plants were photographed at 40 days after planting. Note the difference in silique and SAM scale between *clv3-2*, *S->C 9-3* ‘clv’ and *S->C 10-5* ‘clv’ groups.

The *C* transgene was transformed into *clv3-2* homozygous mutant plants in the Ler background to ensure that this construct produced functional CLV3 protein and that the MmuMiR124 insertion did not affect gene function. Fifteen independent lines were recovered with *C* genes with the *MmuMiR124’* site, and all of these lines rescued the *clv3-2* phenotype. Likewise, eight independent *C* transgenic lines lacking the *MmuMiR124’* site all rescued the *clv3-2* mutant in the T1 generation. Most rescued plants produced phenotypically ‘Wt’ flowers, except for a few flowers produced near the transition to reproductive growth stage that produced an occasional extra petal. From these 15 *clv3-2 C* plants, two independent T4 homozygous lines (*C9*, *C10*) were selected for introduction of the *S* gene. Importantly, the *C9* and *C10* lines displayed a stable ‘Wt’ phenotype over many generations (>T5) and individuals (>1000); i.e., in these genotypes no ‘clv’ phenotypes were ever observed.

The *S* gene was then transformed into the *C9* and *C10* lines (see Figure S7 for a pedigree of the experiment) and 28 and 27 independent *S* lines were recovered in the *C9* and *C10* backgrounds, respectively (hereafter *S->C 9* or *S->C 10*). In the T1 generation, other than some minor aberrations in floral bud formation, all lines appeared phenotypically ‘Wt’ (Figure S8). In the T2 generation, 27 *S->C* lines displayed minor qualitative changes in inflorescence buds, including slight changes in petal number and position, but otherwise appeared to be ‘Wt’ (Figure S9). However, three T2 families, *S->C 9-2*, *S->C 9-3,* and *S->C 10-5* produced phenotypically ‘clv’ plants (Figure S9). One other line, *S->C 9-11*, produced enlarged flowers with more and larger petals and sepals. However, since it did not phenocopy the prototypical *clv3* loss of function, it is not discussed here further. Importantly, all independent ‘clv’ T2 plants were kanamycin resistant, indicating that co-suppression between the *S* and *C* drug resistance genes did not occur and could not have caused the emergence of ‘clv’ plants. Once the ‘clv’ phenotype appeared in the *S->C* lines it was inherited in all self-cross progeny. *S->C 9-3* ‘Wt’ plants homozygous for *S* continued to produce both ‘clv’ and ‘Wt’ progeny at a rate near 25% (102/350 T5 plants; p = 0.07; Χ^2^ test vs 0.25), suggesting the ‘clv’ phenotype was controlled by one recessive locus (Figure 4e&f).

The *S->C* ‘clv’ plants were qualitatively less severe *clv3-2* plants, with visibly less undifferentiated tissue in the inflorescence meristem and less severe silique phenotypes (Figure 4). Additionally, *S->C* ‘clv’ plants often produced flowers with tightly closed sepals, whereas this was rarely observed in the *clv3-2* mutant. To quantify the severity of the ‘clv’ phenotype in the *S->C 9-3* and *S->C 10-5* lines, the number of anthers and petals produced per flower were determined (Figure 5). Both *S->C 9-3* and *S->C 10-5* plants displayed a number of anthers similar to the *clv3-2* mutant. However, *clv3-2* plants produced more petals than either *S->C* ‘clv’ group. Therefore, both *S->C* ‘clv’ lines exhibit phenotypes consistent with loss of *clv3* function, but not as severe as the *clv3-2* null.

**Figure 5.**
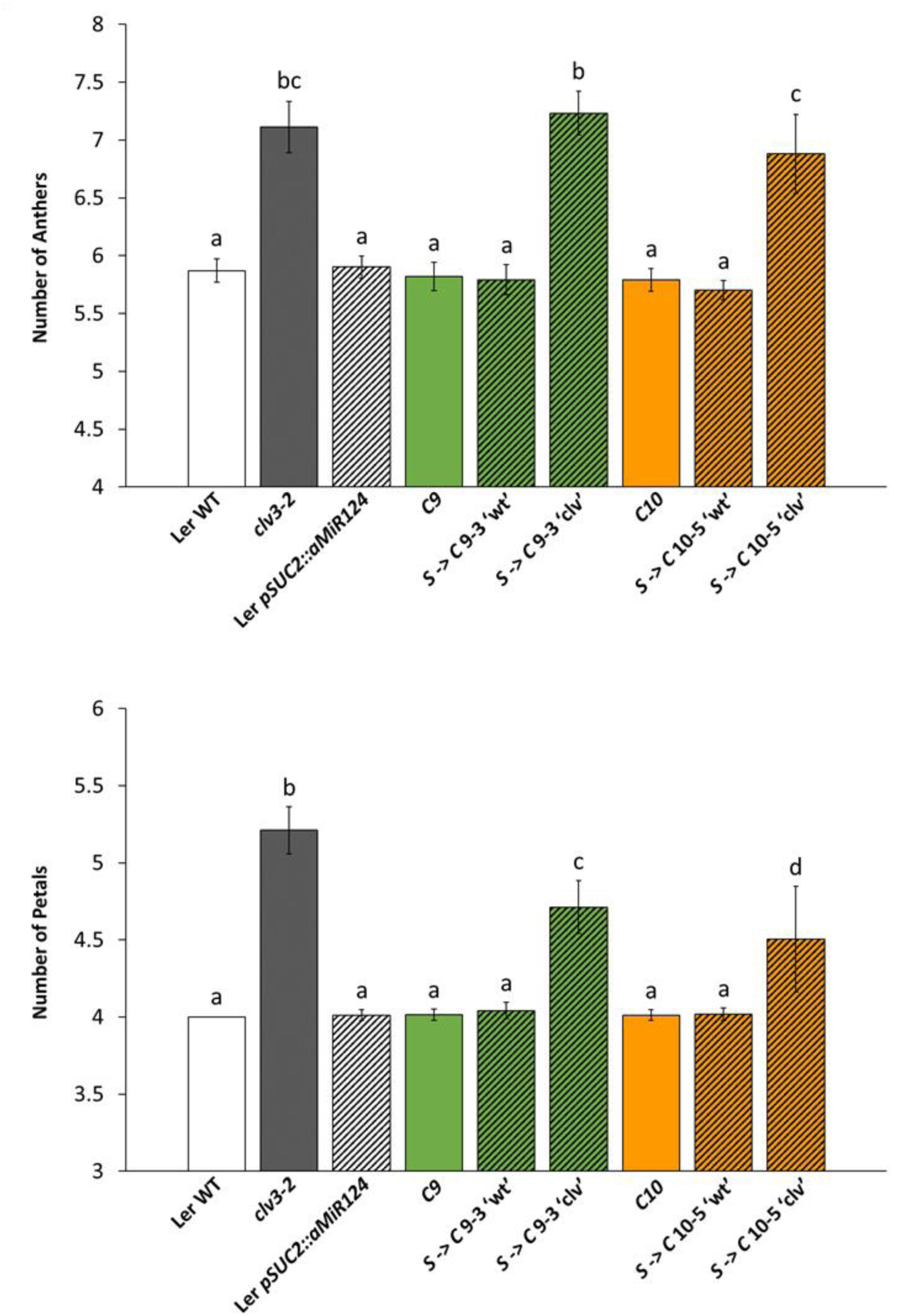
The two independent *S->C* ‘clv’ lines produce additional anthers and petals per flower. All ‘clv’ groups produced aberrant numbers compared to Ler Wt, the *C* only lines, or the *S->C* ‘wt’ groups. Both *S->C 9-3* ‘clv’ *and S->C 10-5* ‘clv’ lines produced a similar number of anthers per flower compared to *clv3-2*. However, *clv3-2* produced more petals per flower than either line. Letters denote statistically similar (p>0.05) groups. For anther counts p values were determined ANOVA with *post hoc* Tukey’s HSD test. For petal counts, due to the zero variance in Ler Wt, p values were determined by a Kruskal-Wallis rank sum test followed by pairwise Wilcox tests. Error bars indicate +/- standard error.

To ensure that the *S* transgene itself does not cause a phenotype resembling loss of *clv3* function, *S* alone was transformed into the Ler Wt background. Of 9 independent *S* lines, none exhibited a *clv*-like phenotype in generations T1 to T4. Likewise, crossing *S* into the *clv3-2* mutant background had no effect on ‘clv’ phenotype. Some *S* lines produced serrated large rosette leaves, which was also seen in the *S->C* lines, including the three lines that produced ‘clv’ individuals (Figure S10). Additionally, these *S* plants had a delayed floral transition, producing more rosette leaves than Ler Wt (Figure S11). The *S* increase in rosette leaf number appeared additive with the ‘clv’ mediated increase in rosette leaf number, with *S->C* ‘clv’ plants producing more rosette leaves than *clv3-2* plants (Figure S12). These observations suggest that, despite the lack of genomic homology, phloem expression of *MmuMiR124* affects Arabidopsis development, but is not responsible for the ‘*clv’* phenotypes in *S->C* lines.

qPCR was used to measure transcript levels of *CLV3* in *S->C* lines that produce both ‘Wt’ and ‘clv’ plants. Both *S->C 9-3* ‘Wt’ and *S->C 10-5* ‘Wt’ plants showed a modest decrease in *CLV3* expression compared to the *C9* and *C10* lines, respectively (Figure 6). This decrease in *CLV3* expression suggests that *S* reduces *C* expression in *S->C* ‘Wt’ plants, but not enough to manifest a ‘clv’ phenotype. The *clv3-2* plants used in this study produced a large amount of transcript, similar to what was reported for the *clv3-9* null allele (Rodriguez-Leal *et al*. 2019). *S->C 9-3* ‘clv’ and *S->C 10-5* ‘clv’ plants also demonstrated an increase in presumably defective *CLV3* transcript (Figure 6). However, consistent with the milder ‘clv’ phenotypes in the *S->C* lines both *S->C* ‘clv’ lines had reduced *CLV3* expression compared to the *clv3-2* background. These data show that *S* derived sRNA may move to decrease *C* expression and, in the *S->C* events that do trigger a ‘clv’ phenotype, the plants over-express non-functional *clv3-2* transcript.

**Figure 6.**
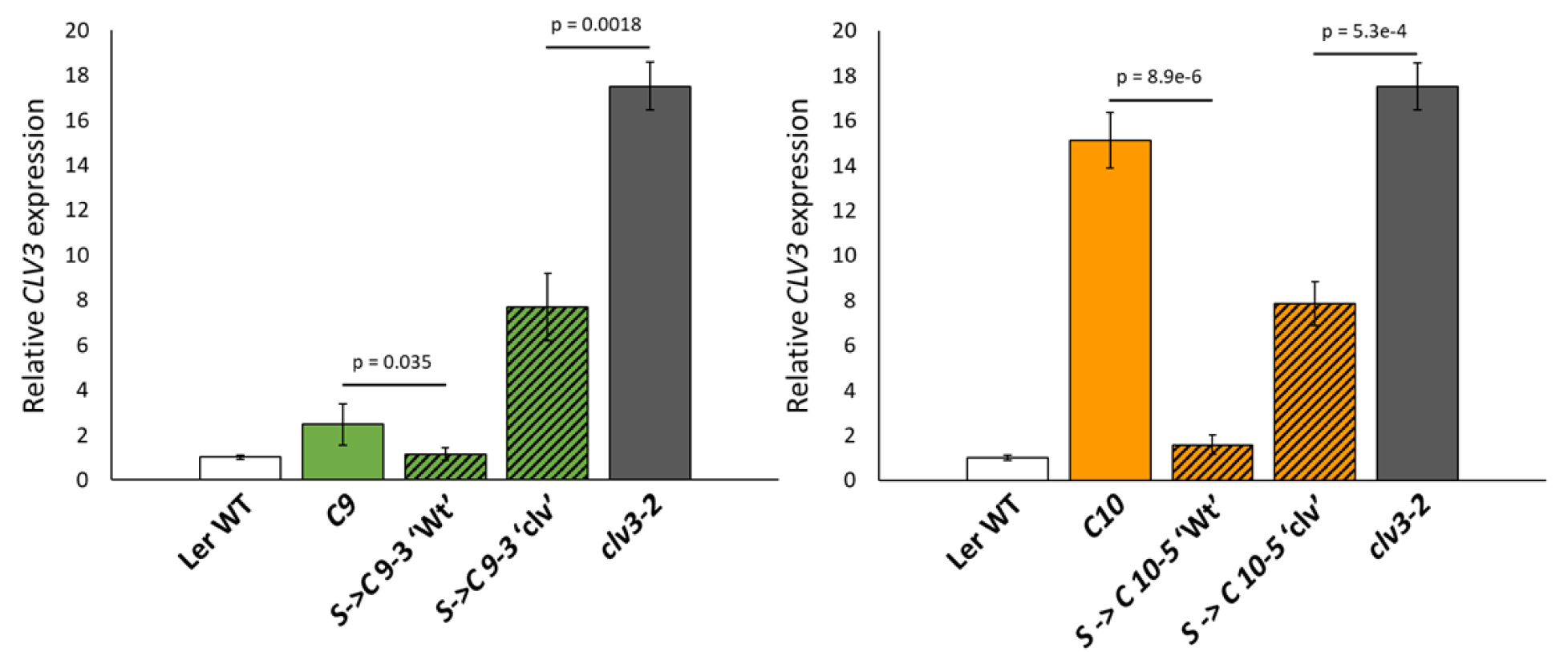
qPCR of apically enriched tissues show differing *CLV3* expression in *S->C* ‘Wt’ and *S->C* ‘clv’ plants. For both independent events, *CLV3* expression was reduced in *S->C* ‘Wt’ plants compared to that observed in their respective *C* only progenitors. Both *S->C 9-3* ‘clv’ and *S->C 10-5* ‘clv’ groups show elevated *CLV3* expression. However, this expression was not as high as that found in the *clv3-2* background. Displayed p values were determined by a student’s t-test. Error bars indicate +/- standard error.

### Strong *S->C* ‘clv’ events appear consistent with epigenetic alterations at *C*

Since *S->C* ‘clv’ events are consistent with the inheritance of a single recessive locus, we investigated the possibility that the *S* gene was no longer required for the ‘clv’ phenotype. That is, is the appearance of the ‘clv’ phenotype in the T2 generation due to permanent silencing of the *C* allele in the T0/T1 generation by *S*? To check this, a *S->C 9-3* ‘clv’ plant was crossed to the homozygous *C9* parental line. If the ‘clv’ phenotype requires continuous *aMiR* silencing, *C9* x *S->C 9-3* ‘clv’ only F2 plants with at least one copy of *S* and two copies of an unlinked recessive *C* allele, should be ‘clv’ (3/16 F2 plants). However, if ‘clv’ is caused by one recessive allele alone, ¼ of the *C9* x *S->C 9-3* ‘clv’ F2 population should be ‘clv’. All F1 plants were ‘Wt’ and 152/528 F2 plants exhibited a ‘clv’ phenotype. This is not consistent with a model that requires the continuous interaction between *S* and *C* (Χ^2^ test vs 3/16; p =3.433e-9). The observed ratio is close to that expected under the single locus model (Χ^2^ test vs ¼; p = 0.0465), consistent with the ‘clv’ phenotype no longer requiring presence of *S.* Furthermore, several *C9* x *S->C 9-3* F2 ‘clv’ plants were genotyped for the presence of the *aMiR* cassette. Of these ‘clv’ plants, 14/64 (Χ^2^ test vs ¼; p = 0.56) lacked *S*. Progeny from four of the F2 lines that genotyped negative for the *aMiR* cassette (which contains a kanamycin resistance gene) were grown on selection and all exhibited kanamycin sensitivity. This again suggests that *S* assorts independently of the ‘clv’ trait. Therefore, *S* likely acted in the T0/T1 generation to silence one *C* allele but is no longer required to maintain the ‘clv’ phenotype.

A possible explanation for the low frequency of *S->C* ‘clv’ events, and their eventual independence from *S*, is that *S* elicited stable epigenetic silencing at *C* in the T0/T1 generation. Alternatively, an unknown mechanism could cause a T0/T1 genetic change producing the *S->C* ‘clv’ phenotype. If such a change caused the *S->C* ‘clv’ phenotype, all plants within the same transgenic line should display an identical ‘clv’ effect. Inconsistent with this genetic explanation, within the same independent line, the severity of the ‘clv’ phenotype changed significantly in subsequent generations (Figure 7). An independent growth experiment comparing two *S->C 9-3* T4 families again supported ‘clv’ phenotype differences within the same transgenic lines (Figure S13). This variation in phenotype is at odds with a genetic explanation of the *S->C* ‘clv’ events, suggesting T0/T1 epigenetic changes may be involved.

**Figure 7.**
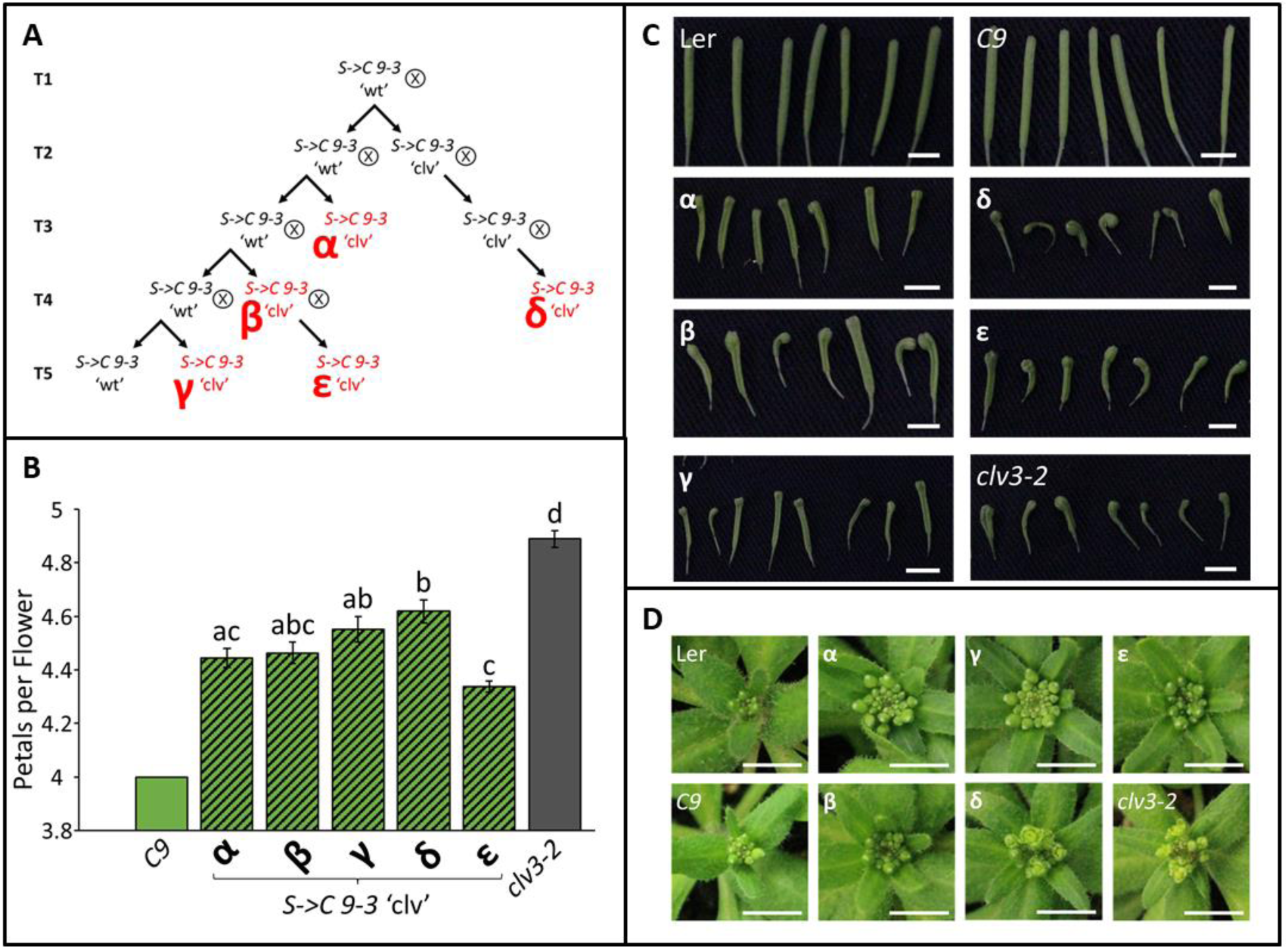
*S->C* ‘clv’ families exhibit phenotypic variation between genetically identical plants. (**A**) A pedigree outlining the relatedness of the sampled *S->C 9-3* families. Sampled families are written in red and identified by a Greek letter (α-ε). All families were homozygous for both S and C. (**B**) The number of petals produced per flower varied within different *S->C 9-3* ‘clv’ families.’ (**C**) Silique and () developing inflorescence phenotypes produced by the *S->C 9-3* transgenic families. Mirroring the petal number data, with the δ family exhibiting the most severe qualitative ‘clv’ phenotypes. **D** Letters denote statistically similar (p>0.05) groups, as determined by ANOVA with post hoc Tukey’s HSD test. C9 was excluded from this statistical analysis to allow the use of a parametric test. Error bars indicate +/- standard error. Scale bars denote 0.5 cm.

### *S->C* phenotypes exhibit somatic instability and Treatment of S->C ‘clv’ plants with 5-azacytidine produced one whole plant reversion

Close observation of *S->C* individuals revealed occasional somatic instability, with ‘clv’ plants producing ∼1-3 siliques that appeared phenotypically ‘Wt’ (Figure 8; Figure S14). ‘Wt’ siliques were never observed in *clv3-2* mutant plants (Figure S14). Seed produced from ‘Wt’ siliques (on an otherwise ‘clv’ plant) were sown to check for changes in progeny phenotype. Two out of 288 plants from these ‘Wt’ siliques produced phenotypically ‘Wt’ plants, suggesting these somatic sectors can represent heritable reversions. Conversely, although rarer, *S->C* ‘Wt’ plants could produce ‘clv’ siliques with 3-4 carpels, suggesting both *C* alleles had been silenced in a sector (Figure 8). When seed from ‘clv’ siliques on an otherwise ‘Wt’ plant were grown, they segregated ‘Wt’ plants only, indicating the sectored silencing was not heritable. Taken together, these observations reveal somatic instability of both the ‘Wt’ and ‘clv’ phenotypes in the *S->C* background. To investigate the possibility that the *C* locus in ‘clv’ individuals was methylated, ‘clv’ plants were treated with the DNA methylation inhibitor 5-Azacytidine (5-Aza). One of the 719 F3 seeds treated with 5-Aza produced a ‘Wt’ plant. The progeny of this 5-Aza revertant plant segregated ‘clv’ as a recessive trait, consistent with one *C* allele having regained function (Χ^2^ test vs ¼; p = 0.84 n = 34). The ‘Wt’ progeny of this 5-Aza revertant displayed flowers with unclosed sepals, which was not observed in *C9* or *S->C 9-3* ‘Wt’ lines but is seen ‘clv’ individuals. This is consistent with the 5-Aza treatment reactivating one *C* allele, although not to the same level as the naïve state before *S* introduction.

**Figure 8.**
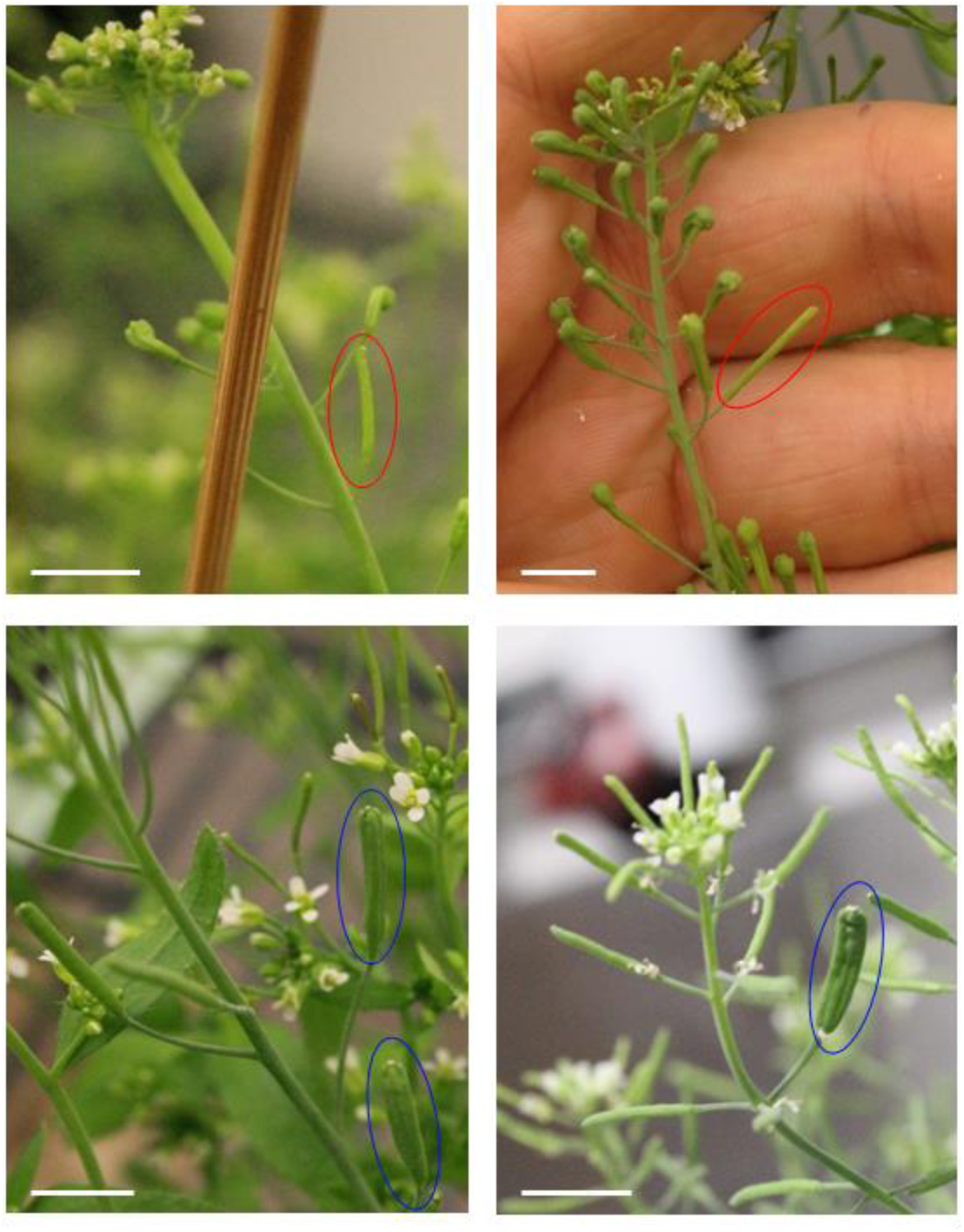
*S->C* plants exhibit somatically unstable phenotypes, consistent with revertant or silenced sectors. *S->C* ‘clv’ plants can produce one or two bi-carpellate ‘Wt’ siliques (top, red ovals). Likewise, *S->C* ‘Wt’ plants can produce ‘clv’ multi-carpellate siliques (bottom, blue ovals). Scale bar denotes 1 cm.

### *S->C* plants display variable shoot morphologies that are not heritable

*S->C* transgenic plants exhibited occasional shoot aberrations. Examples include inflorescences that end in a terminal flower or plants that lose apical dominance (Figure S15). Paradoxically, similar bud termination effects have been observed in plants that overexpress *CLV3,* suggesting an imbalance of CLV3 signaling in *S->C* plants (Müller *et al*. 2006). This phenotypic variability was more pronounced after the plants were outcrossed. In three *S->C* F2 populations, a wide array of unexpected polymorphisms appeared (Figure S16). None of these phenotypes were heritable, although the phenotypic instability continued into the F3 generation. F2 populations resulting from Ler x *S* displayed no phenotypic instability, suggesting the instability required both *S* and *C*. One commonly observed phenotype was inflorescences that terminated with multi-carpellate or multi-staminate structure, reminiscent of flowers in *carpel factor*y (aka *dcl1*) mutants (Jacobsen *et al*. 1999). These variable growth aberrations suggest that interaction of *S* and *C* might stochastically disrupt shoot growth and development, possibly by interfering with endogenous sRNA signaling.

## Discussion

### Phloem-derived sRNA can influence gene expression in the SAM

The mobility of sRNA within the plant has been established, yet the extent of this movement with respect to phloem-to-SAM remains unclear (Voinnet *et al*. 1998; Brosnan and Voinnet 2011). Systemic transitive silencing can reach the SAM, but this represents the culmination of a series of short distance events, not long-distance trafficking between distant regions of the plant. Importantly, phloem-to-SAM sRNA trafficking allows for distally generated sRNA signals to reach the SAM, even if they cannot induce systemic transitivity, allowing the SAM to be influenced by diverse sRNA signals originating in remote parts of the plant. The transgenic systems described here suggest that an axis of communication from phloem-to-SAM via sRNA messages does exist, and provides preliminary insights into how endogenous phloem-to-SAM sRNA signaling occurs.

In both synthetic messenger systems described here the sRNA signal originates in phloem companion cells and migrates to different targets. The *pSUC2::FDi* system demonstrated that fully complementary inverted repeat RNA expressed in phloem could downregulate gene expression in the SAM. Curiously, *FD* repression was lost in the *rdr6-15* mutant. RDR6 catalyzes the creation of dsRNA from ssRNA precursors, a step that is not needed to produce sRNA from inverted repeats, like *FDi* (Himber et al. 2003; Peragine et al. 2004; Allen et al. 2005). Therefore, *rdr6-15* must disrupt long distance silencing downstream of the initial sRNA biogenesis. Inverted repeats generate a spectrum of sRNA sizes that can trigger secondary sRNA biogenesis and transitivity, and we suspect this is likely what is lost in the *rdr6-15* background (Bleys et al. 2006; Wroblewski et al. 2014). Importantly, *FD* is expressed exclusively in the apex (Abe *et al*. 2005; Wigge *et al*. 2005). Therefore, secondary sRNA biosynthesis cannot be needed for sRNA to reach the SAM, as it could only occur after the sRNA has reached *FD*. Thus, RDR6-mediated transitivity is likely a critical step in phloem-to-SAM sRNA signal reception, amplifying a weak mobile signal to elicit gene silencing. miRNAs can sometime trigger secondary sRNA biogenesis, and it is possible the *S->C* system likewise relies on RDR6 actions in the SAM (Allen *et al*. 2005; Creasey *et al*. 2014). In this way, only small initial populations of phloem mobile sRNA need to reach the SAM to trigger an effective silencing response.

### sRNA signals can target the stem cells of the shoot apex

The *S->C* silencing system combined a phloem-expressed *aMiR* with a synthetic *CLV3* target expressed in SAM stem cells. Most *S->C* transgenics were phenotypically ‘Wt,’ but showed reduced *CLV3* transcript levels supporting phloem-to-SAM trafficking of *S*-derived miRNA to the central zone. Only a few independent lines produced ‘clv’ individuals, which is consistent with strong enough knockdown of *CLV3* to manifest meristem over-proliferation (Reddy and Meyerowitz 2005; Müller et al. 2006). The notion that epigenetic events underly the *S->C* ‘clv’ events was supported by the appearance of revertant somatic sectors and significant variation phenotype within the same transgenic line.

Study of miRNA intra-SAM movement describe distinct cellular domains wherein miRNAs act non-autonomously; outside of these domains, non-autonomous miRNA silencing was restricted (Skopelitis *et al*. 2018). In contrast, our *S->C* system supports miRNA movement from the phloem through these domain barriers to stem cells in the central zone. In most instances (*S->C* ‘Wt’ plants), *S*-triggered reduction in *CLV3* expression was modest, perhaps below levels that can be detected by the histological approach used in Skopelitis *et al*., (2018). Additionally, the transgenics described in Skopelitis *et al*., (2018) were in an *rdr6^-^* background, which may preclude strong silencing events in the T0/T1 generation (see below). Taken together, we suggest that sRNA gating within the SAM may not be impregnable, such that some dosage of miRNA may pass through these domain barriers. Indeed, a similarly leaky sRNA gating mechanism was observed in the parenchyma subtending the SAM (Liang *et al*. 2012).

### Stochastic T0/T1 events may allow for strong *C* silencing events

The appearance of *S->C* ‘clv’ in the T2 generation suggests that only one *C* allele was silenced in the T0/T1 generation. Therefore, we speculate that initial *trans*-actions of *S* triggered *cis* silencing at one allele. The rarity of such events (3/55 insertions; 3/110 *C* alleles), suggest some stochastic trigger for strong *C* silencing. The *C* aMiR site is close to the transcriptional terminator, the disruption of which has been associated with sRNA-mediated transgene silencing (Brand et al. 2002; de Felippes et al. 2020). If any *S*-derived sRNA initiates RdDM, it may spread DNA methylation onto the linked terminator sequence, silencing *C* (Ahmed et al. 2011). miRNAs have been demonstrated to elicit RdDM, often by triggering transitivity, and we suspect this entry into RdDM could be the stochastic event behind strong *C* silencing. In *Nicotiana benthamiana*, *miR319a* produces 21nt miRNAs, which are not typically associated with transitivity (Chen et al. 2010). However, the *aMiR* has an altered miRNA sequence, and miRNA structure can cause 22nt miRNA production and transitivity (Iki et al. 2018). Therefore, if *S*-derived 22nt sRNA can reach the central zone they could silence expression from the *C* transgene. Alternatively, both *AGO7* (transitivity-associated) and *AGO6* (RdDM-associated) are expressed in the central zone and could interact with *S*-derived sRNA to silence *C* (Montgomery et al. 2008; Tian et al. 2014; McCue et al. 2015; De Felippes et al. 2017).

We suspect that all strong *C* silencing events occurred in the T0/T1 generation, as no ‘clv’ events later emerged from transgenic lines that did not show T2 ‘clv’ phenotypes. The T0/T1 generation is critical for sRNA silencing of newly inserted transgenic transposons, with T0/T1 sRNA silencing mutants having transgenerational effects on the efficacy of silencing (Fultz and Slotkin 2017). Since strong *C* silencing occurred shortly after *S* integration, it is possible that integration transiently affects sRNA silencing of not only the integrated sequence, but also of any homologous loci. T-DNA transgenes come from a bacterial genome and they lack cytosine methylation and any pre-existing histone state. Perhaps events during *de novo* establishment of a chromatin state during *S* integration enables the plant genome to occasionally facilitate RdDM at *C*. DNA integration involves dsDNA breaks, which alone induce sRNA expression (diRNA) and are thought to recruit RNA Pol IV and V (Wei et al. 2012; Miki et al. 2017). How dsDNA break triggered events affect *de novo* chromatin modifications remain unclear, but perhaps they provide a brief window wherein *S* silencing was capable of strong *C* silencing. Importantly, diRNA synthesis requires RNA Pol II transcription, so any *S* diRNA would be co-expressed with *pSUC2* and, if they are needed to silence *C*, still require phloem-to-SAM trafficking (Miki et al. 2017). Future investigations should study how the molecular events associated with integration influence sRNA silencing in *cis* and *trans*.

### Long distance sRNA trafficking could induce epigenetic changes at the SAM

The role of sRNA in mediating gene silencing is well established. Therefore, phloem-to-SAM sRNA transport provides a route for somatic sRNA to regulate genes in the stem cells that produce the shoot, gametes and, consequently, the next generation. Mobile sRNA can stably alter DNA methylation in tissues, including the root apical meristem, and we suggest similar events occur in the SAM (Melnyk *et al*. 2011a; Lewsey *et al*. 2016; Yu *et al*. 2018). It is intriguing to speculate that, since plants do not completely erase DNA methylation during gametogenesis, epigenetic changes caused by phloem-mobile sRNA to be inherited (Quadrana and Colot 2016). The composition of sRNA populations in the phloem change with the environment, raising the possibility that environmentally induced sRNA could have lasting epigenetic consequences, both in this generation and the next (Buhtz et al. 2008; Pant et al. 2008). Indeed, others have suggested that sRNA could bridge environmental sensing with progeny imprinting, and our demonstration of phloem-to-SAM sRNA transport provides a route for these adaptive epigenetic changes to occur (Martienssen 2008, 2010; Brosnan and Voinnet 2011; Minow and Colasanti 2020).

## Methods

### Plant materials and growth conditions

*Arabidopsis thaliana* plants were grown in Sunshine mix LA4 or Promix BX at 60% relative humidity, with an irradiance of 150 µmolm^-2^s^-1^ and constant day/night temperatures of 22°C and 18°C respectively. Plants were fertilized bi-weekly with liquid 17-5-17 (200ppm) fertilizer. Plants were grown under 16-hour long days photoperiods, or 8-hour short day photoperiods when specified. Experiments were conducted in the Landsberg *erecta* (Ler) or Columbia-0 (Col) accessions. Arabidopsis mutants (*clv3-2*, *fd-5*, *hen1-6*, *hst-6*, *hst-15*, and *rdr6-15*) were obtained from the Arabidopsis Biological Resource Center (ABRC). Under desired circumstances, hand crosses were conducted using unopened female flowers and dehiscent anthers

### Recombinant plasmid construction

To construct the *pSUC2::FDi* and *pFD::FDi* transgenes, a 264bp region of the first exon of *FD* was amplified from cDNA and inserted into the *pHANN* vector in sense and anti-sense orientation. Then this hairpin-forming fragment was amplified and added to *pDONR221 P5P2* via Gateway® recombination. The *pSUC2* promoter sequence was amplified from a plasmid graciously provided by George Coupland and recombined into *pDONR221 P1P5r*. The *pFDi* promoter was amplified off an Arabidopsis BAC clone (F4B14 from the ABRC) and likewise added to *pDONR221 P1P5r*. A 3-way Gateway® reaction recombined promoters with *FDi* into the pK7WG vector.

The *pSUC2::GUS:GFP* reporter construct was created by Gateway® recombination cloning *pSUC2* into the *pDONR P4P1r* vector, then recombining this into *pKGWFS7*. To construct *C*, the native *pCLV3::CLV3::eCLV3* locus was amplified off Col Wt genomic DNA in two fragments and subcloned into *pUC19* through restriction enzyme cloning. Within the 3’ *CLV3::eCLV3* fragment, a naturally occurring NsiI site (New England Biolabs) was used to insert an oligonucleotide duplex containing the *MmuMiR124’* site into the CLV3 3’ UTR. Next the two *pCLV3::CLV3::eCLV3* fragments were digested and inserted into *pBM42GW,3* via a three fragment ligation. A rescue control was likewise constructed using 3’ *pCLV3::CLV3::eCLV3* without the *MmuMiR124’* insertion. To construct *S*, p*SUC2* inserted into *pK7m24GW,3* via restriction enzyme cloning. Gene synthesis (Eurofins-Operon) was used to create the *aMiR* by replacing the active *miRNA319a* site with that of *MmuMiR124*. This *aMiR* was then cloned into *pK7m24GW,3* downstream of *pSUC2*. Primers used and the final sequences of all expression vectors are contained in Table S1 and Figure S28 respectively. All PCR reactions were performed in a Bio-Rad T100 thermal-cycler using Phusion High-Fidelity DNA Polymerase (Thermo-fisher) or KOD Hot Start DNA Polymerase (Sigma-Aldrich).

### Stable transformation of Arabidopsis

Agrobacterium-mediated Arabidopsis floral dips were used to integrate transgenes as previously described (Clough and Bent 1998), using Agrobacterium strain GV3101. For herbicide selection, plants were grown on ½MS plates containing 35 µgmL^-1^ kanamycin or ½MS plates containing 20 µgmL^-1^ Basta (Glufosinate-ammonium) and 1% (w/v) sucrose. Alternatively, soil grown plants were sprayed three times with 200µgmL^-1^ Basta. Unless otherwise indicated, analysis was conducted on plants homozygous for a given transgene. The *SUC2::FDi* and *FD::FDi* constructs were created in Ler and Col ecotypes, however most quantitative analysis was conducted in the Col background to allow comparison to the *fd-5* mutant. To construct the *S->C* system, *clv3-2* mutants were dipped with the *C* transgene; then, two stable homozygous lines were transformed with *S* (See Figure S7 for a pedigree of the experiment).

### Flowering time and floral organ phenotypic analyses

For quantitative flowering time analysis, Arabidopsis seeds were stratified (3-4 days at 4°C) prior to sowing and thinned to uniformity. For each growth experiment, comparisons were made within the same chamber with all genotypes randomly distributed. Flowering time was determined as the number of days before floral buds were first discerned in the rosette, and rosette leaf number includes all true leaves except those initiated from axillary meristems. To score floral organ number, the number of petals or anthers per flower were counted for nine flowers from plants at the same developmental stage. Individual flower organ counts were averaged to create floral organ counts for individuals. Since floral petal number varied the most between *S->C 9-3* ‘clv,’ *S->C 10-5* ‘clv’ and *clv3-2* plants, only petals were counted from different lineages within the *S->C 9-3* background. For *S->C* ‘clv’ segregation ratios, seedlings were germinated on ½MS plates containing 1% (w/v) sucrose before transplantation to soil, to avoid bias for seedling vigor during thinning. Seeds germinated on plates were first sterilized through sequential washes of 0.05% Triton-X in 70% ethanol, and two changes of 100% ethanol. Rosette leaf area was calculated via ImageJ. All stats were conducted using R (V3.4.1).

### DNA extraction and genotyping

Following *SUC2:FDi* crosses, PCR genotyping for the transgene or *fd-5* T-DNA insertion was conducted using the primers and conditions listed in Table S2. Plants homozygous for *hen1-6*, *hst-6*, *hst-15*, or *rdr6-15* were identified via phenotype. PCR genotyping was conducted using GoTaq® Green Master Mix (Promega) in a Bio-Rad T100 thermal cycler. To extract genomic DNA, leaf tissue was frozen in liquid nitrogen, ground to a powder and resuspended in buffer consisting of 200mM Tris-HCL, 250 mm NaCl, 25mM EDTA and 0.5% (w/v) SDS (pH 7.5). The supernatant was removed, and DNA was subsequently precipitated with isopropanol, pelleted, and washed twice with 70% ethanol. This genomic DNA was then resuspended in water.

### RNA extraction and qPCR expression analysis

The apical regions of 15-day old Arabidopsis seedlings with all mature rosette leaves and roots removed, but including developing leaves and some petiole were immediately flash frozen in liquid nitrogen. This apically enriched tissue was pooled from five plants to produce an independent replicate. RNA was extracted with TRIzol (Invitrogen), per manufacturer instructions. RNA was treated with DNAse I (Thermo-fisher) and subsequently cleaned up via phenol/chloroform extraction and ethanol precipitation. RNA quality was determined via agarose gel electrophoresis and RNA purity and quantity was measured via NanoDrop 2000C (Thermo Scientific). 1000µg total RNA was used in iScript cDNA Synthesis (Bio-Rad). After confirming primer specificity and efficiency via a standard curve, qPCR measurement of transcript abundance was done using SsoAdvanced Universal SYBR Green Supermix (Bio-Rad) on an Applied Biosystems 7300 real-time PCR system. qPCR primers are listed in Table S3. For all qPCR experiments, 3 technical replicates and 3-6 biological replicates were used. To normalize expression to the amount of meristem derived cDNA in the total cDNA pool, *KNAT1* expression was used as a meristem-specific endogenous control in addition to β*-tubulin*. Importantly, *KNAT1* expression showed no expression trends within the groups sampled, implying its expression was unchanged across the genotypes sampled and serves as a good measure of meristem content (Figure S29). The 2^−ΔΔCt^ method was used to determine expression across groups (Livak and Schmittgen 2001). Due to the mild difference in *FD* expression, this qPCR experiment was repeated 3 times independently, all of which exhibited the same trend in *FD* expression.

### 5-Azacytidine treatments

Arabidopsis seeds were sterilized (see plant growth conditions) and plated in sterile conditions upon ½MS plates with or without 100µM 5-Aza, which has been previously demonstrated reduce genome-wide DNA methylation (Griffin *et al*. 2016). Seeds were dark stratified for 3 days at 4°C before being transferred to light. After 8 days of growth all seedlings were transferred to soil and grown to maturity.

### Sectioning, GUS staining and microscopy

Whole seedlings were cleared in 70% ethanol at room temperature over several days. GUS staining was carried out as previously described (Jefferson *et al*. 1987). GUS-stained tissues were counter stained with Eosin Y and embedded in paraffin wax before being sectioned (Leica RM2665) into 8µm ribbons. These ribbons were placed on poly-l-lysine coated slides, re-hydrated, and viewed via light microscopy (Leica DMLS2). GFP fluorescence was viewed via epifluorescence microscopy (Leica MZFLIII). For imaging of Arabidopsis SAMs, the inflorescence tissue was hand dissected before being placed fresh into an environmental scanning electron microscope (Hitachi TM-1000).

## Acknowledgements

Research supported by the Natural Sciences and Engineering Research Council of Canada. MAAM is a recipient of an NSERC doctoral fellowship. We thank Mike Mucci, Tannis Slimmon and Leane Illman of the Guelph Phytotron for expert plant care. We would also like to thank Jeff Gross from the Guelph genomics facility for his service and several excellent conversations.

## Supplemental Figures

**Figure S1.**
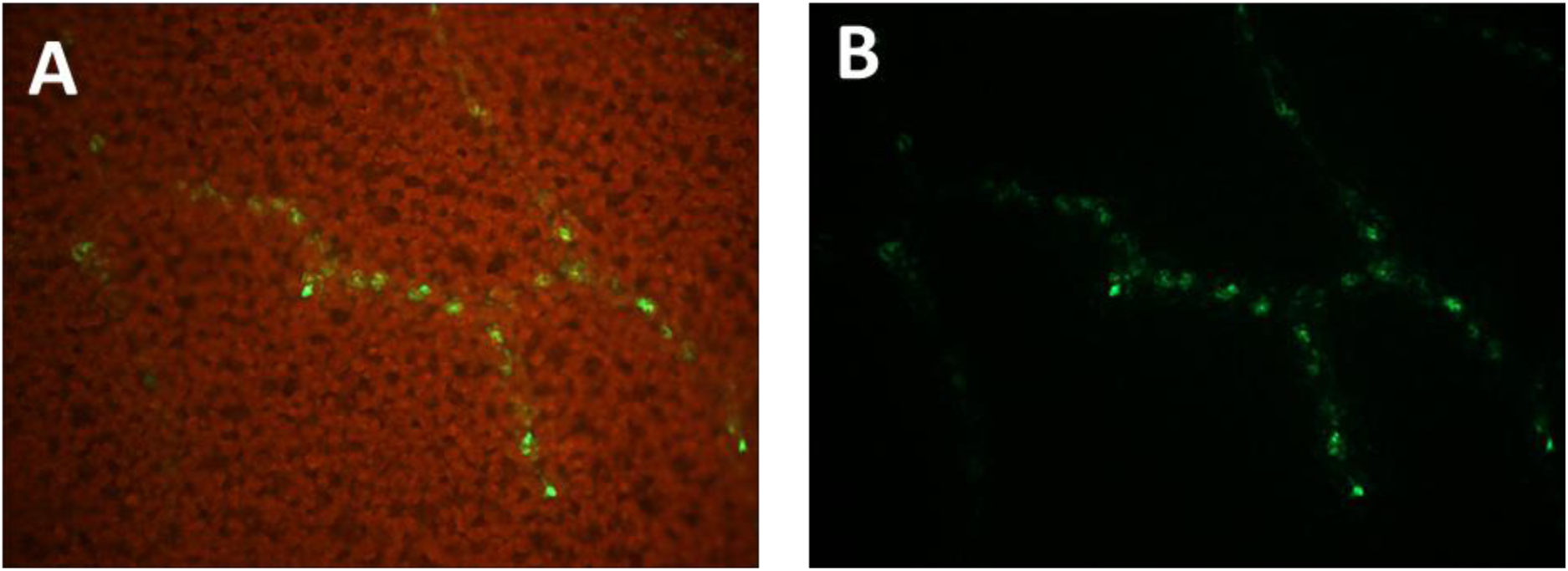
GFP fluorescence pattern from the *pSUC2::GUS:GFP* reporter construct. GFP fluorescence was likewise constricted to the vasculature as with (**A**) or without (**B**) chlorophyll autofluorescence.

**Figure S2.**
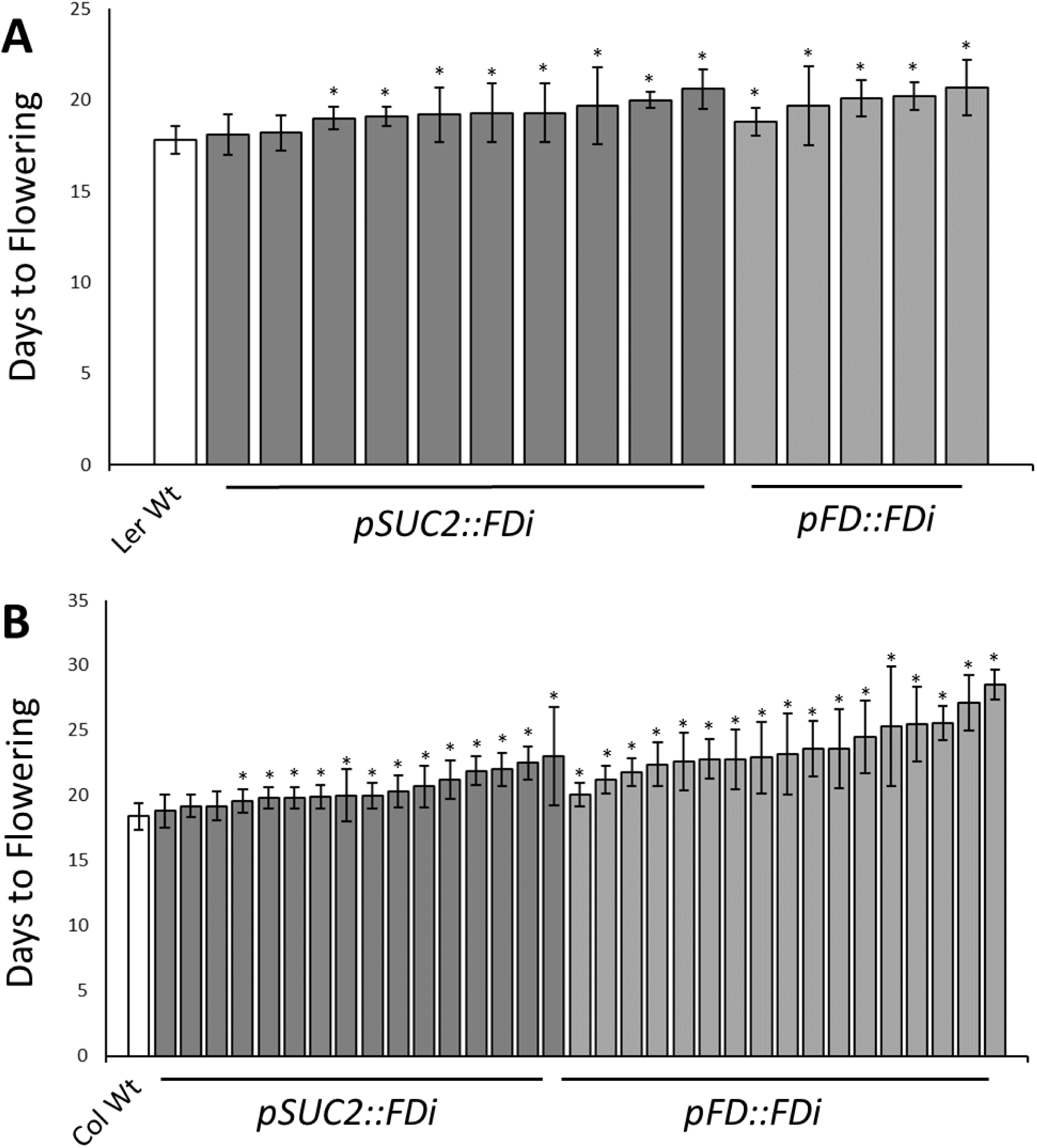
Under LD photoperiods, T2 *pSUC2::FDi* and *pFD::FDi* lines exhibited delayed flowering in both (A) Ler and (B) Col backgrounds. Each bar represents the flowering time of an independent insertion event. T2 plants were confirmed to contain the Kanamycin resistance marker. Significant (p>0.05; t-test) differences between transgenic families and their respective wild types are denoted by a *. Error bars represent +/- standard deviation.

**Figure S3.**
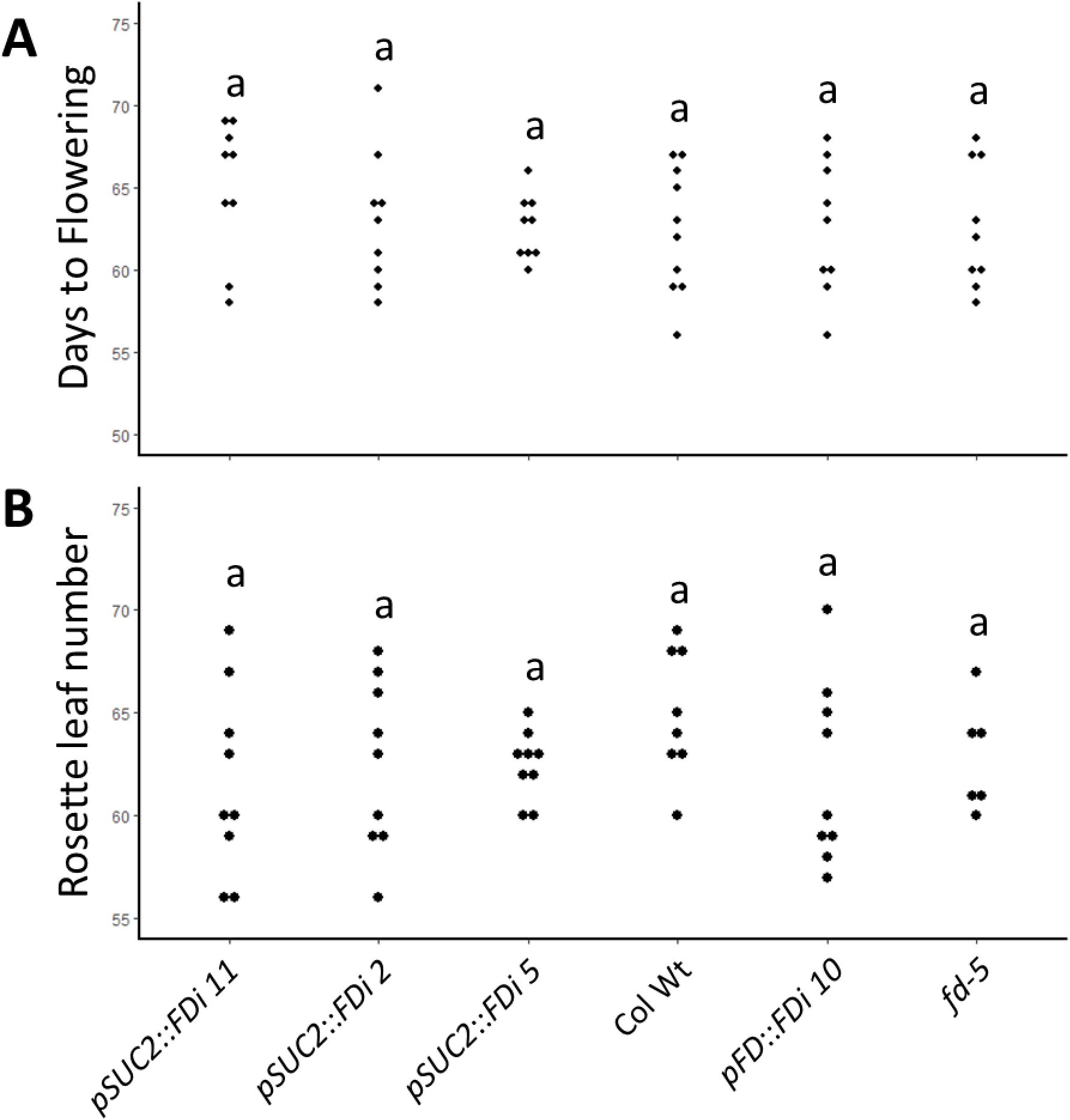
Under SD photoperiods, *pSUC2::Fdi, pFD::FDi* and *fd-5* lines exhibited flowering time equivalent to Col Wt. (**A**) All genotypes flowered at a similar day after planting and (**B**) produced an equivalent number of rosette leaves. Each dot represents an individual. Letters denote statistically similar (p>0.05) groups, as determined by ANOVA with post hoc Tukey’s HSD test.

**Figure S4.**
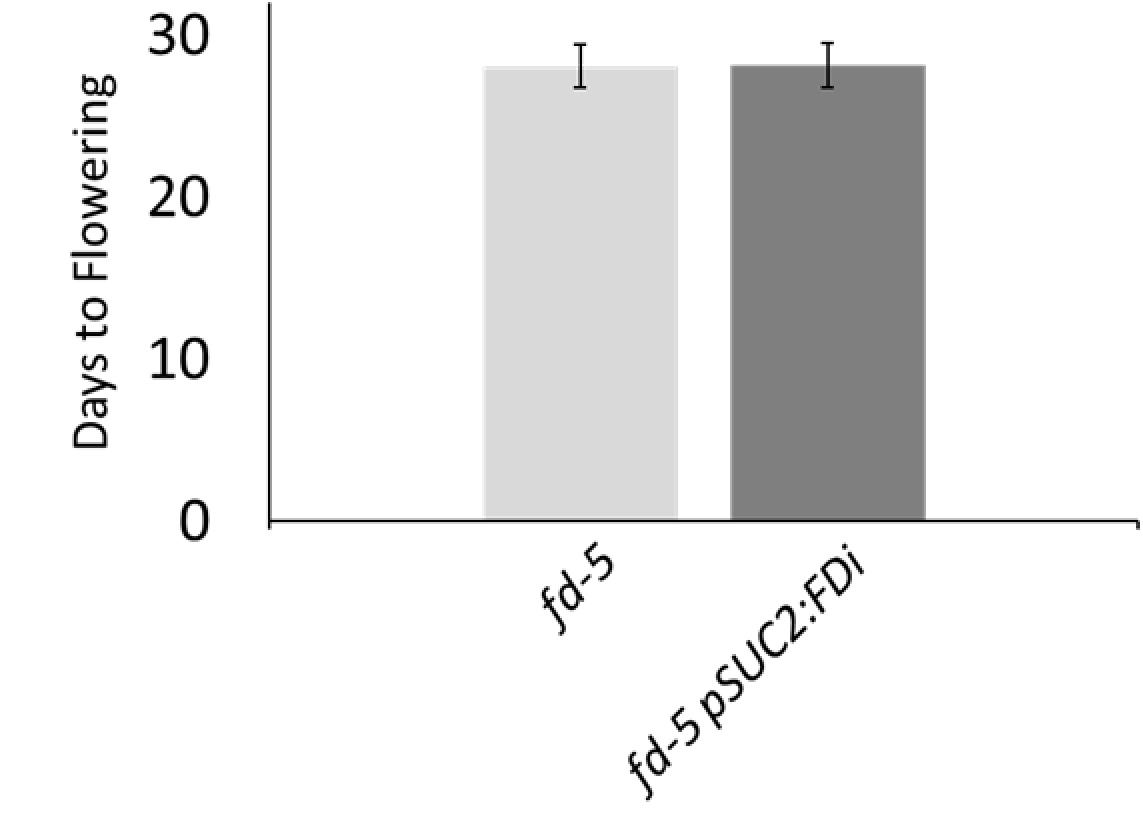
*pSUC2::FDi* has no additive effect on flowering when in a *fd-5 loss* of function background. Both *fd-5* and *fd-5 pSUC2::FDi* had statistically equivalent (p=0.85; t-test) days to flowering when grown together under long day conditions. Error bars denote +/- standard deviation.

**Figure S5.**
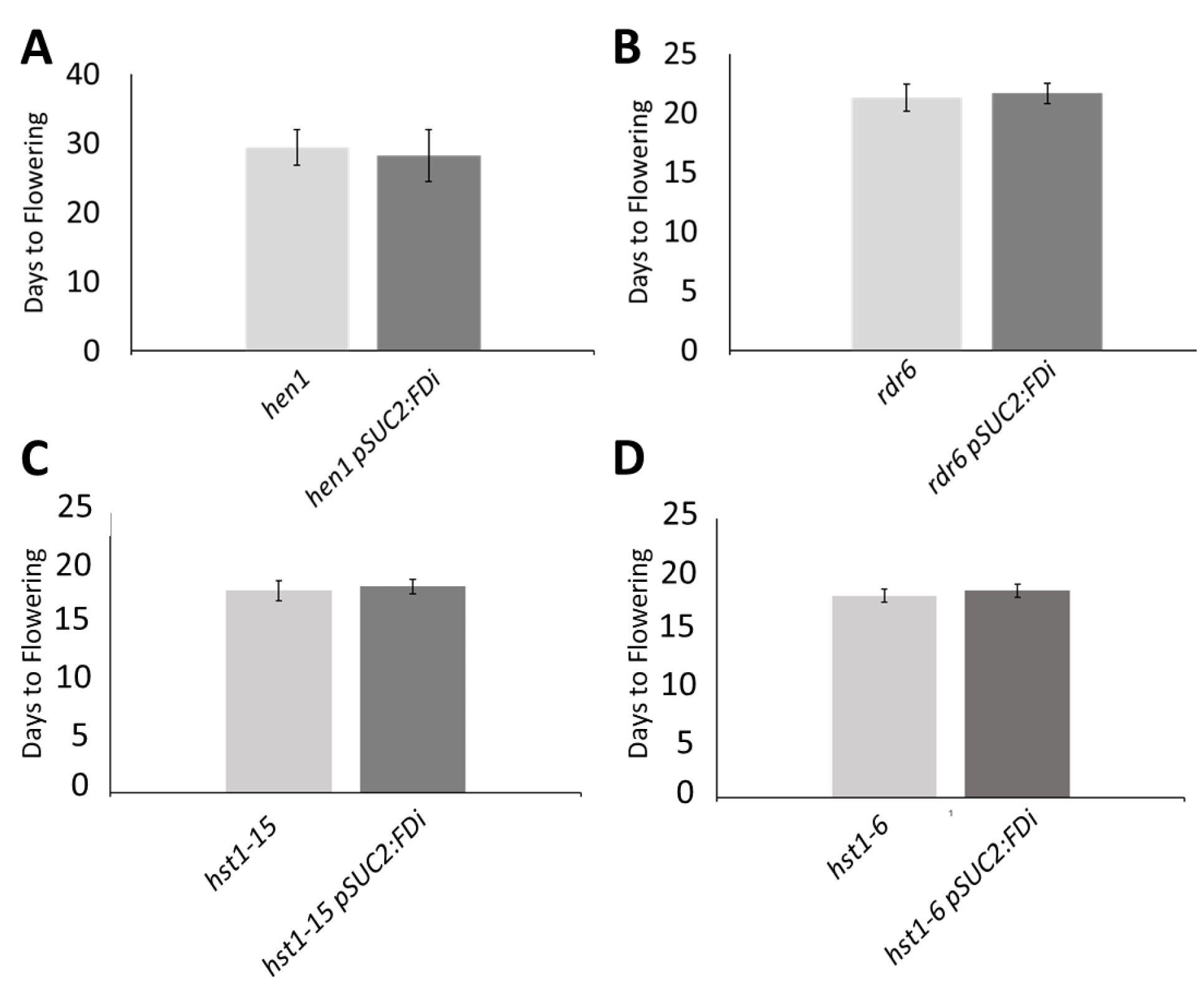
The floral delays caused by *pSUC2::FDi* is disrupted when combined with known RNAi mutants. The *pSUC2::FDi* floral delay was abolished when combined with (**A**) *hen1*, (**B**) *rdr6*, or (**C&D**) two *hst* alleles. Pairwise comparisons within all mutant backgrounds illustrated that presence of the *pSUC2::FDi* transgene had no significant (p>0.05;t-test) effect on days to flowering. Error bars denote +/- standard deviation.

**Figure S6.**
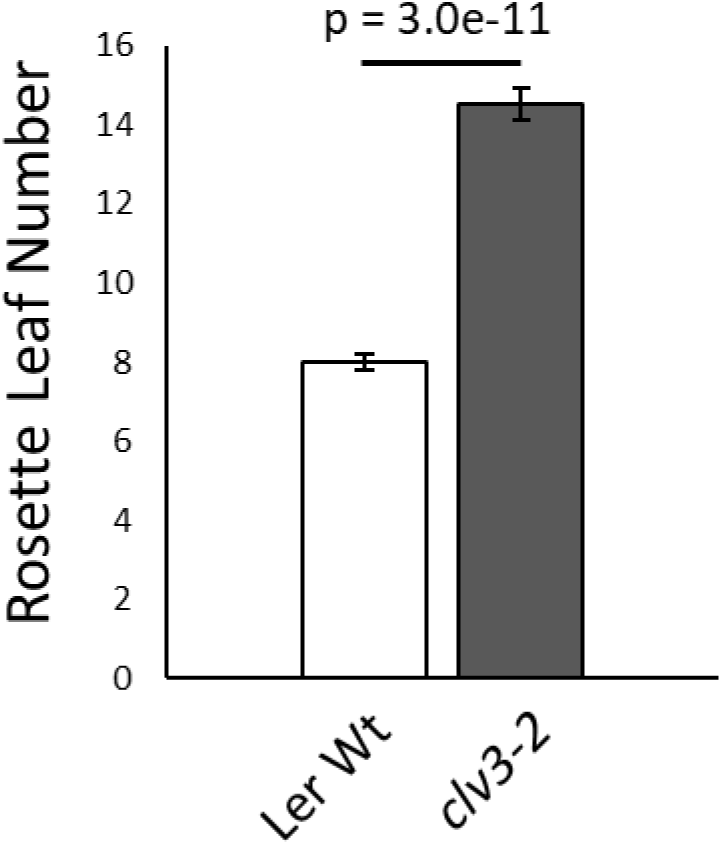
Loss of function *clv3-2* mutants produce more rosette leaves before flowering. The p value of a pair-wise t-test is displayed. Error bars denote +/- standard error.

**Figure S7.**
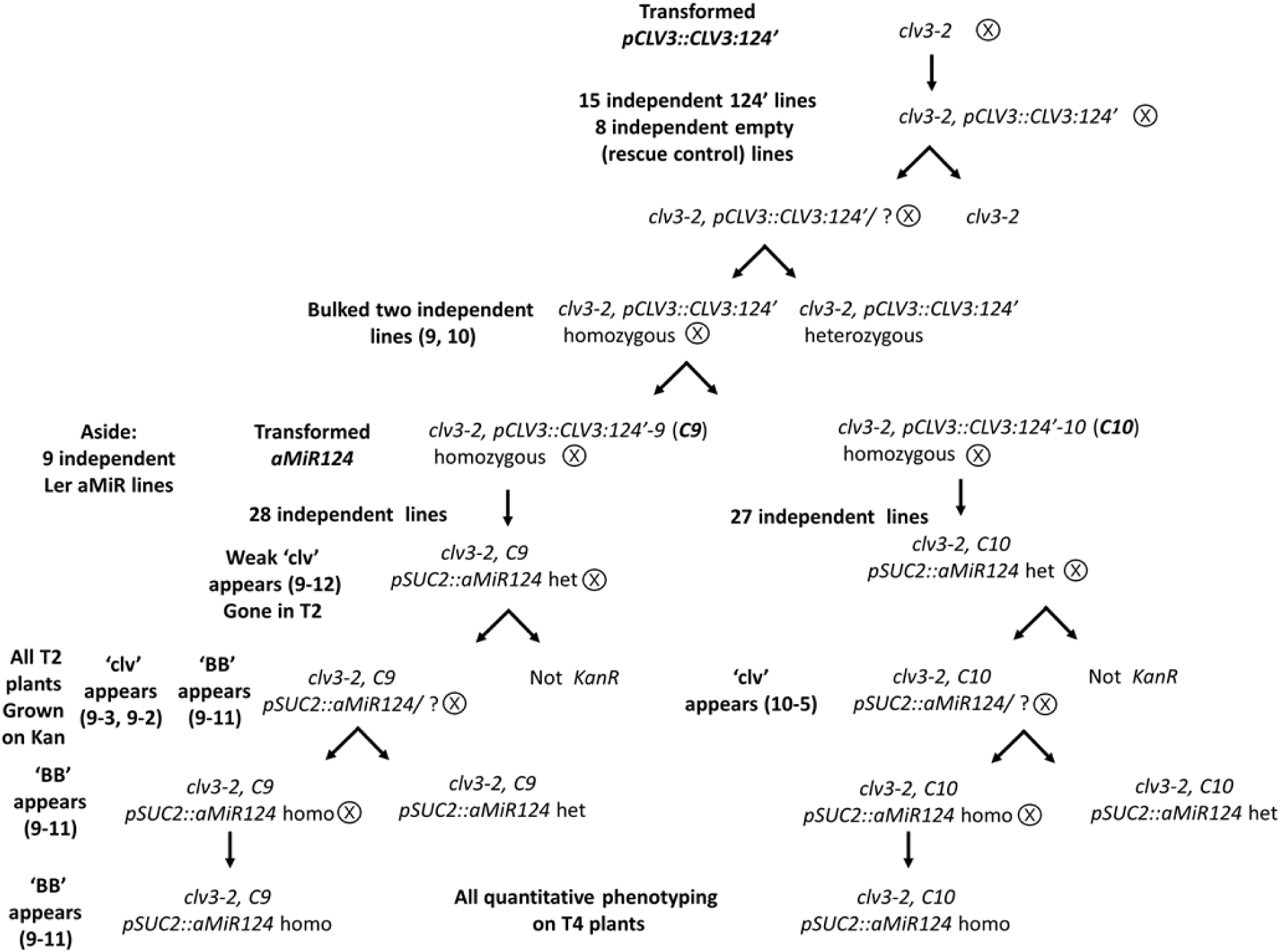
*S->C* experiment pedigree. First, *clv3-2* mutants were transformed with the *pCLV3::CLV3:124*’ (abbreviated to *C*) transgene as well as a rescue control of the *C* cassette lacking the *MmuMiR124’* site. As expected, Eight rescue controls and all 15 *C* lines rescued the *clv3-2* phenotype. From these *C* lines, two independent lines (*C9* & *C10*) were selected on the basis of phenotype stability. *pSUC2::aMiR124 (*abbreviated to S) was then transformed into these lines, as well as into the Ler background alone. 28 and 27 independent lines were recovered in the *C9* and *C10* backgrounds respectively. During the generation of homozygous lines, ‘clv’ plants first appeared in the T2 generation. The big bud (‘BB;’ see text for description) also appeared in the T2, but continued to segregate in more advanced generation. Quantitative comparison between lines first took place on T4 homozygous lines.

**Figure S8.**
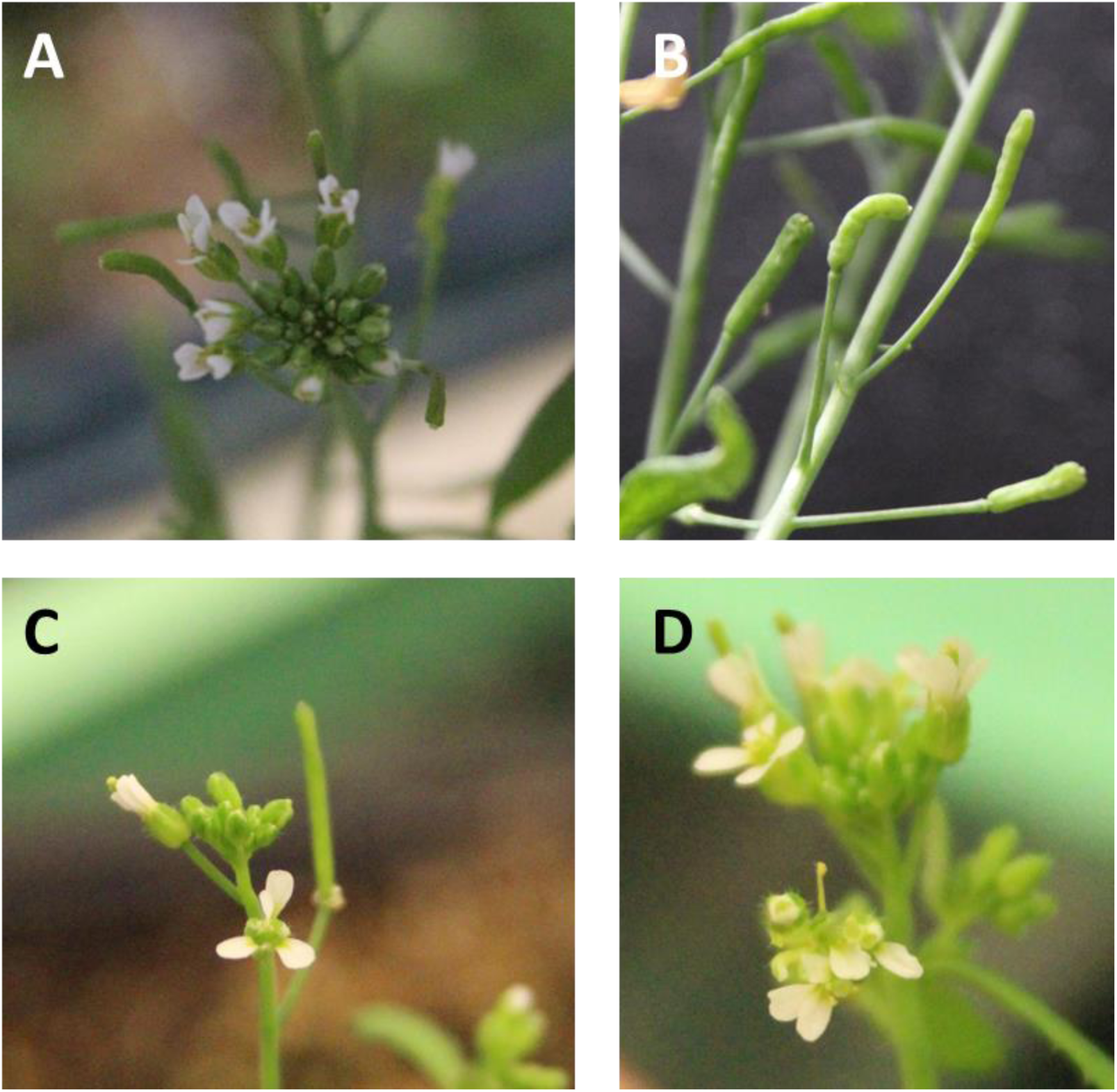
Representative images of the minor shoot changed observed in independent T1 *S->C* lines. *S->C* T1 plants often displayed (**A&B**) mishappen siliques, (**C**) aberrant petal arrangement or (**D**) termination of inflorescences into a determinate floral structure.

**Figure S9.**
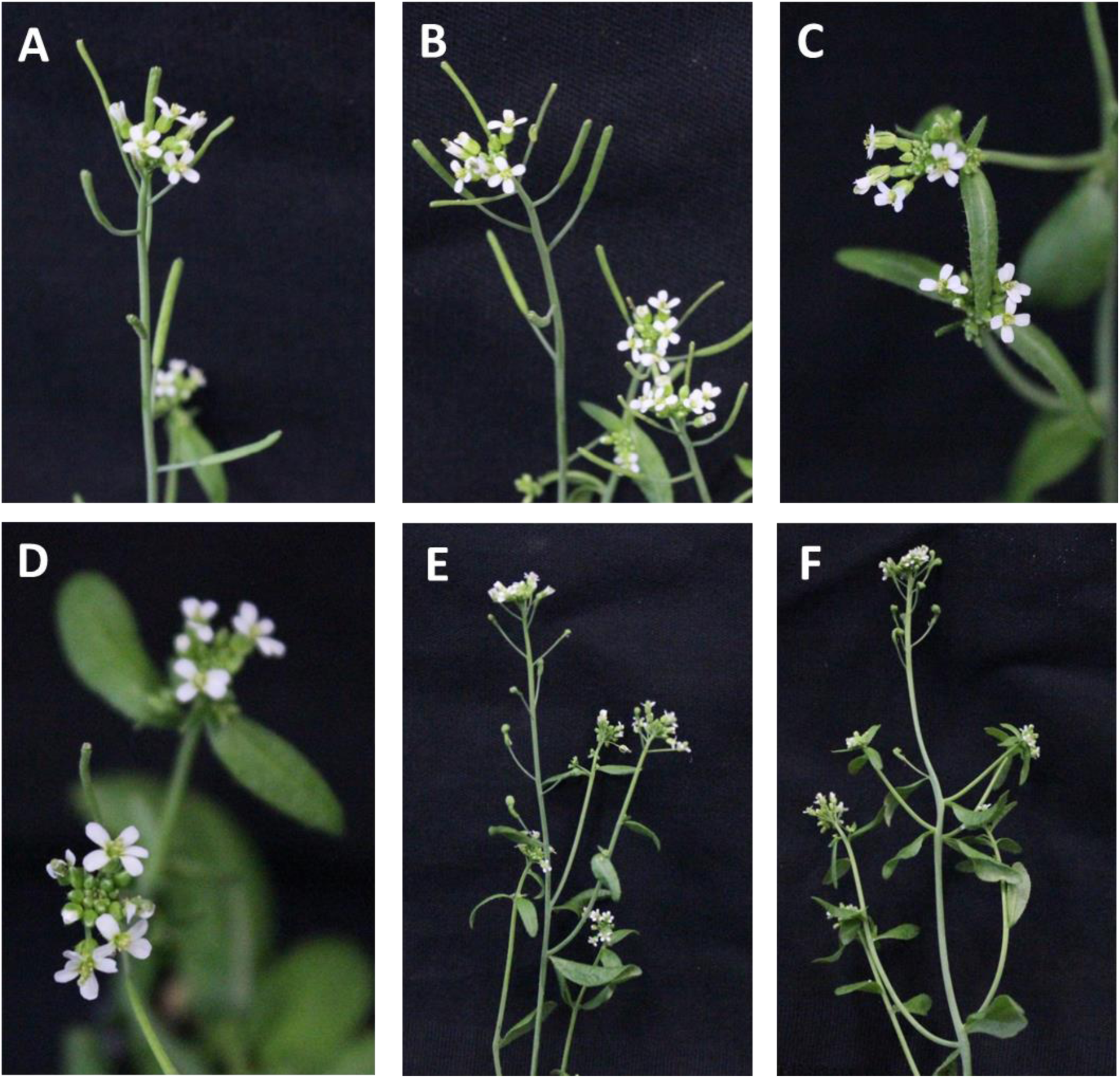
Representative images of the shoot changed observed in independent T2 *S->C* lines. The (**A**) *C10* or (**B**) *C9* rescue lines produced plants that are phenotypically very close the ‘Wt’. (**C&D**) Most *S->C* lines produced a very similar ‘Wt’ phenotype, only occasionally producing flowers with extra petals. However, (**E**) *S->C 9-3* and (**F**) *S->C 10-5* lines produced strong ‘clv’ individuals. One other line, *S->C 9-2*, was later found to likewise produce ‘clv’ individuals (not depicted here). All plants displayed contained the Kanamycin resistance marker lined to *S*.

**Figure S10.**
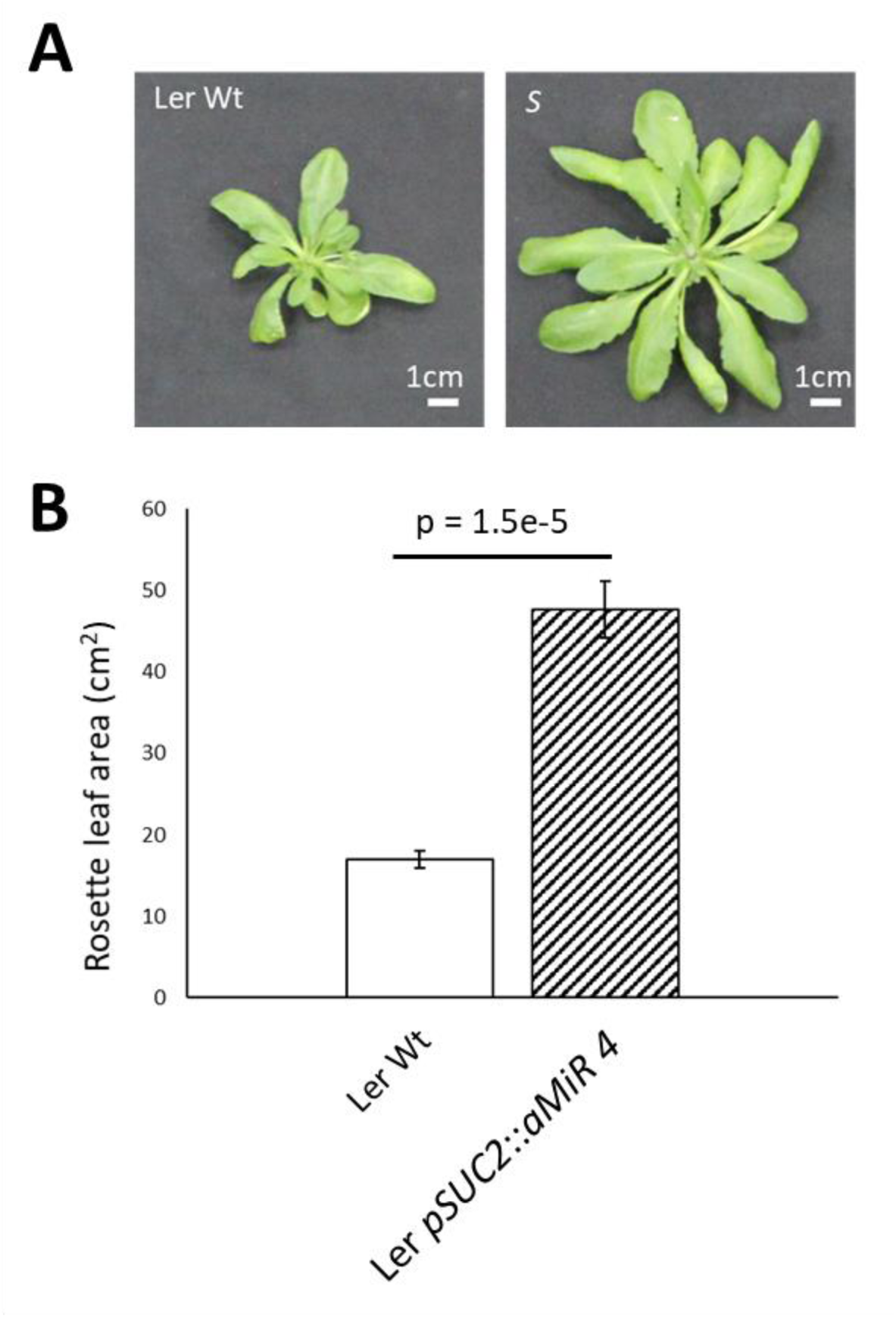
Transformation of *S* into Ler Wt produced larger and serrated rosette leaves. (**A**) Representative image of a 35-day old Ler Wt (Left) and an *S* line in the Ler background (right). (**B**) *S* transgenics produced larger rosette areas when compared. A similar change in rosette leaves was seen in *S->C* lines after the transformation of *S*. The p value of a pair-wise t-test is displayed. Error bars denote +/- standard error.

**Figure S11.**
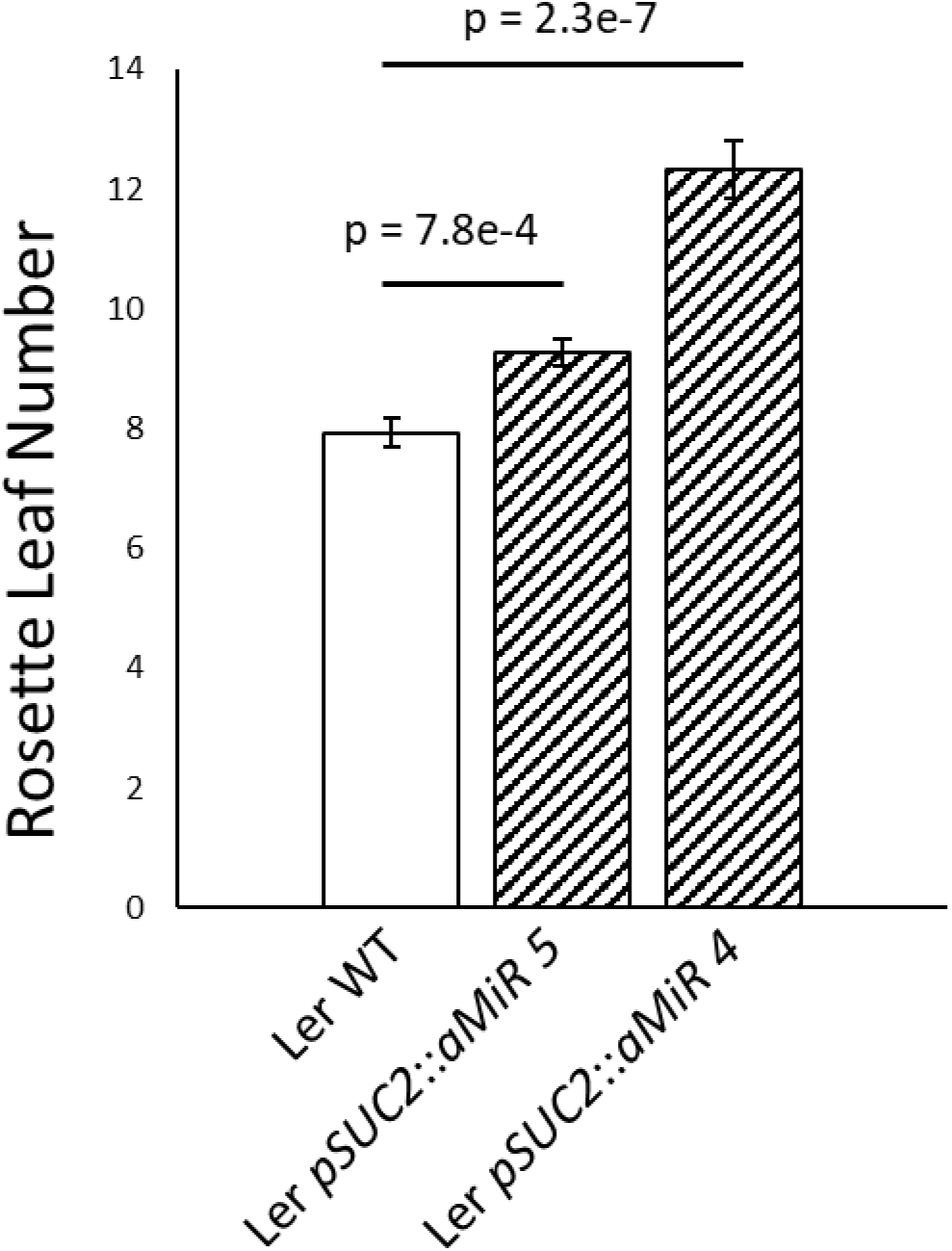
*S* plants produced more rosette leaves than Ler Wt. A similar change in rosette leaf number was seen in *S->C* lines after the transformation of *S*. The p values of pair-wise t-tests are displayed. Error bars denote +/- standard error.

**Figure S12.**
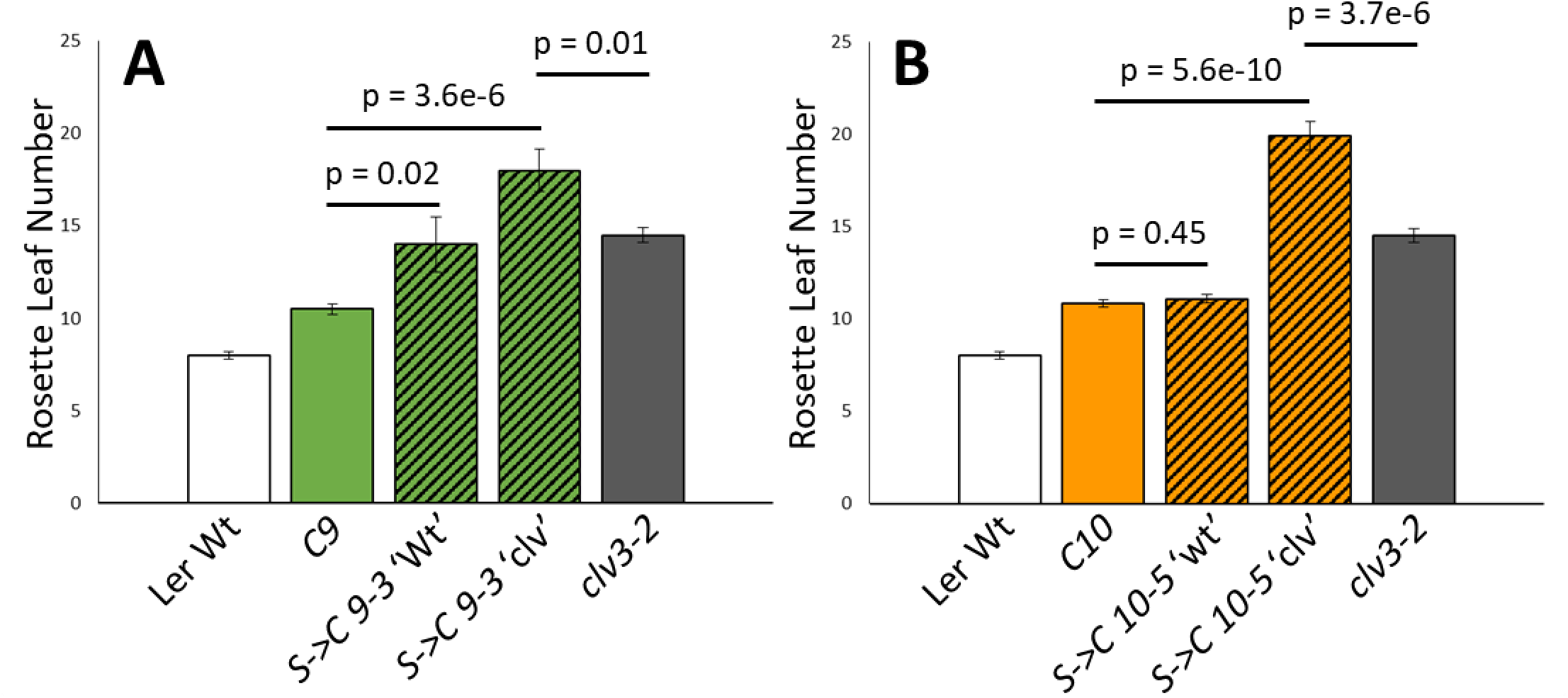
The increase in rosette leaf number caused by *S* and ‘clv’ appeared additive in *S->C* plants. (**A**) *S->C 9-3* ‘Wt’ plants produced more leaves than *C9*, and *S->C 9-3* ‘clv’ plants produced more leaves than *C9* or *clv3-2.* (**B**) *S->C 10-5* ‘clv’ plants produced more leaves than *C10* or *clv3-2.* However, *S->C 10-5* ‘Wt’ plants did not produce more leaves than *C10* in this grow out, consistent with *S* variably impacting rosette leaf number. The p values of pair-wise t-tests are displayed. Error bars denote +/- standard error.

**Figure S13.**
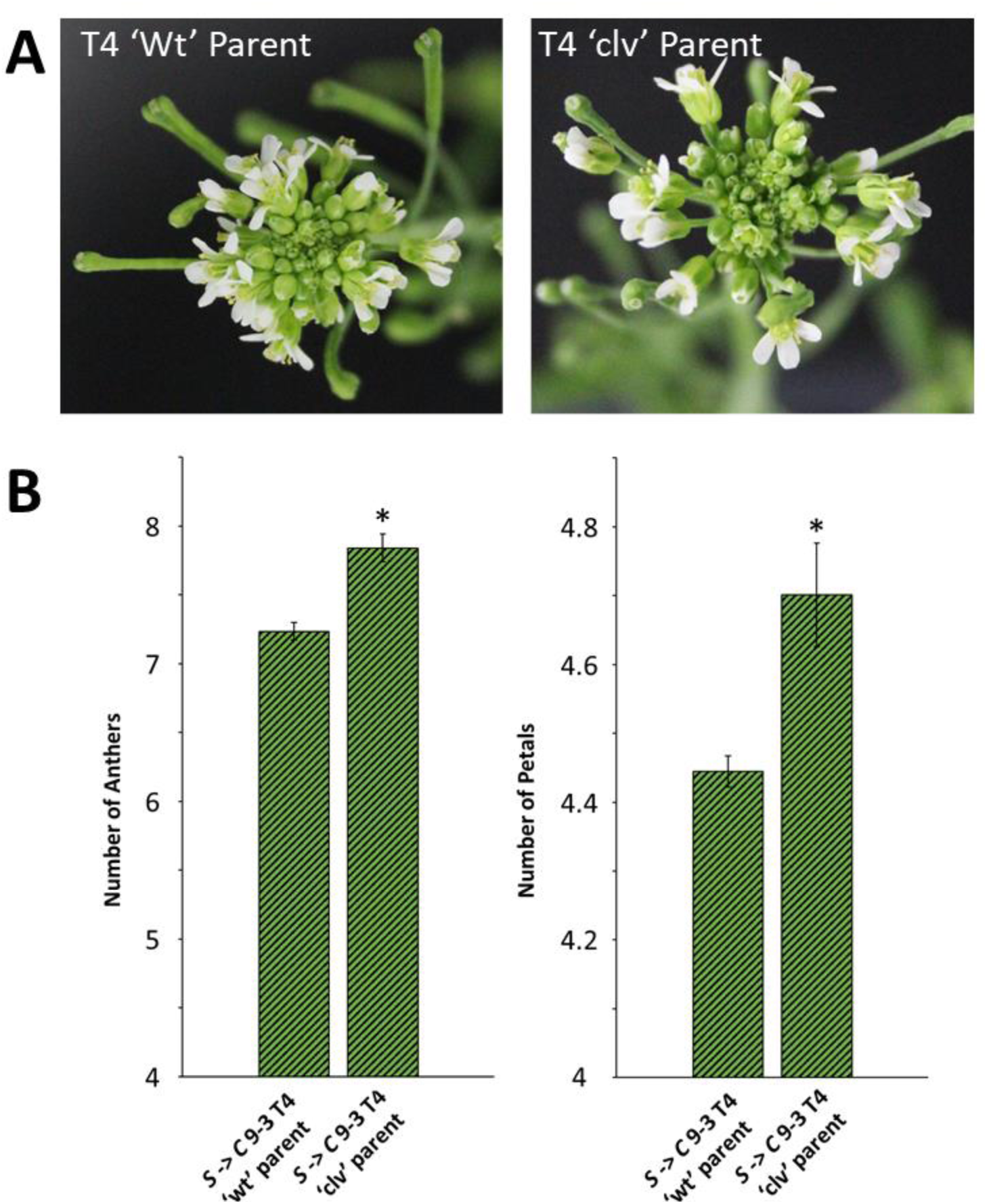
*S->C 9-3* transgenics consistently produce intra-family differences ‘clv’ phenotype severity. (**A**) A second comparison of *S->C 9-3* families confirmed the qualitative differences in silique and apex morphology seen previously. (**B**) Pairwise comparison of the number of petals and anthers produced per flower was consistent with the *S->C 9-3* T4 family from a ‘clv’ parent was more severe than that produced from a separate ‘Wt’ parent. All plants are homozygous for both *S* and *C*. Significant (p>0.05; t-test) differences between families are denoted by a *. Error bars represent +/- standard error.

**Figure S14.**
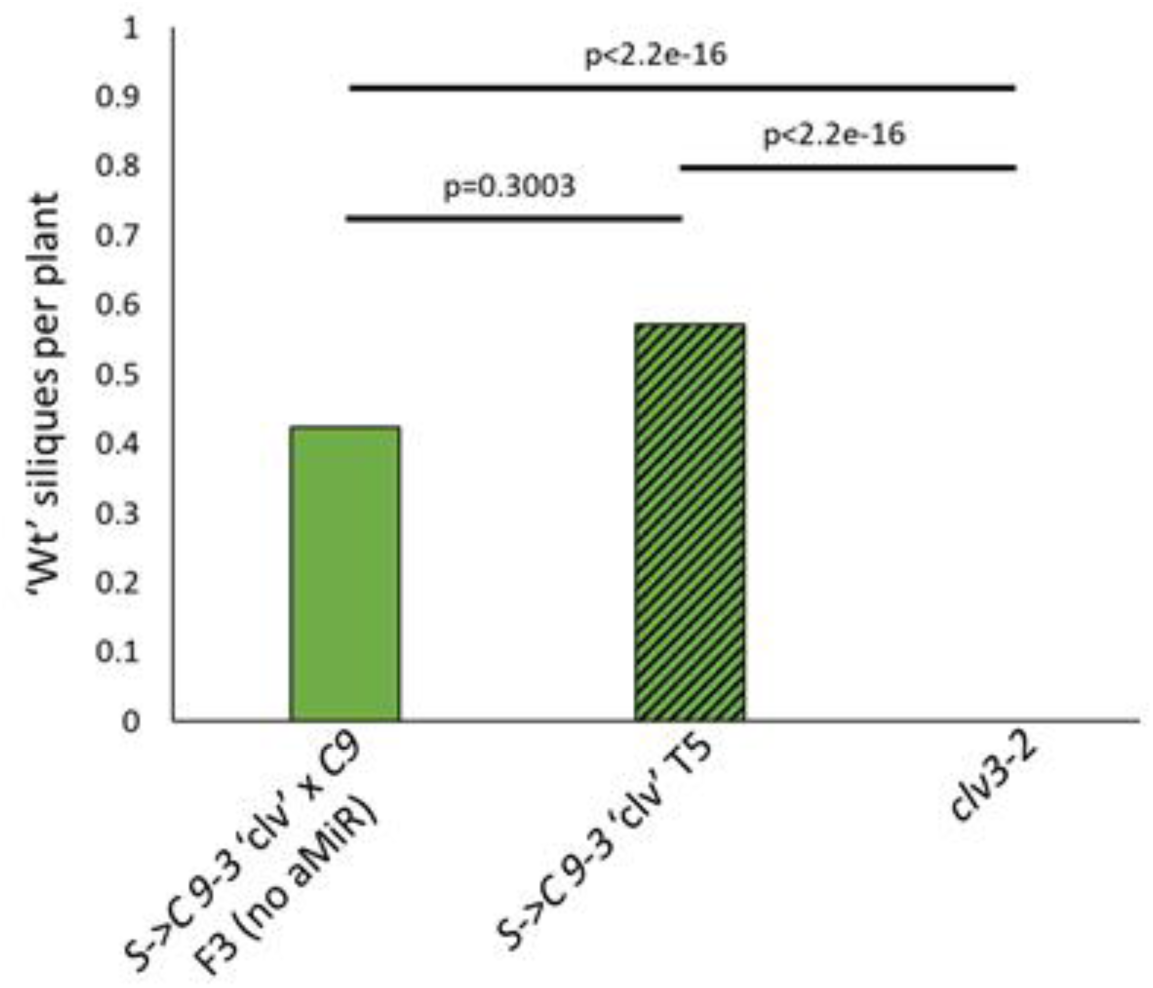
No ‘Wt’ siliques were seen on *clv3-2* plants and the number of ‘Wt’ siliques per plant was similar between related plants, regardless of the presence of *S*. The p values of exact Poisson tests are displayed (n values are 33, 35 and 33 for *S->C 9-3* ‘clv’ X *C9* F3, *S->C 9-3* ‘clv’ T5, and *clv3-2* respectively)

**Figure S15.**
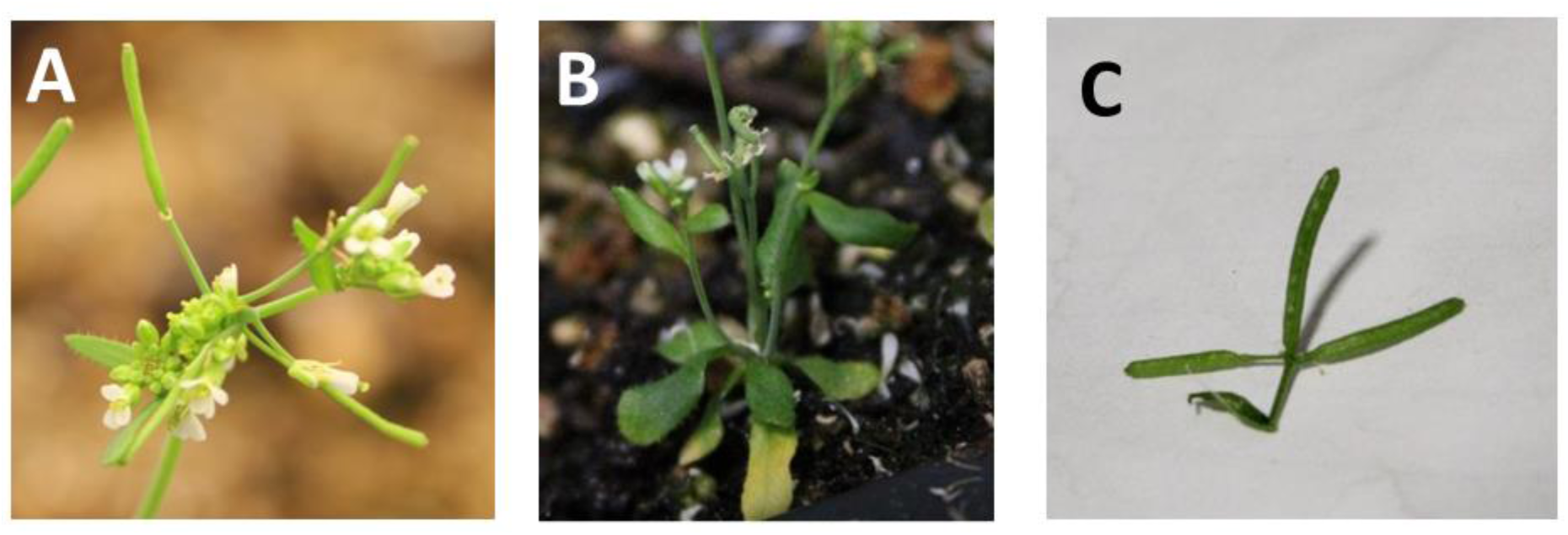
*S->C* plants can produce terminal inflorescences. These inflorescences were observed in various lines including (**A**) *S->C 9* T1 (**B**) *S->C 9-11* ‘Wt’ X *S->C 9-11* ‘BB’ F2 and (**C**) *S->C 9-3* ‘clv’ X *C9* F3 plants.

**Figure S16.**
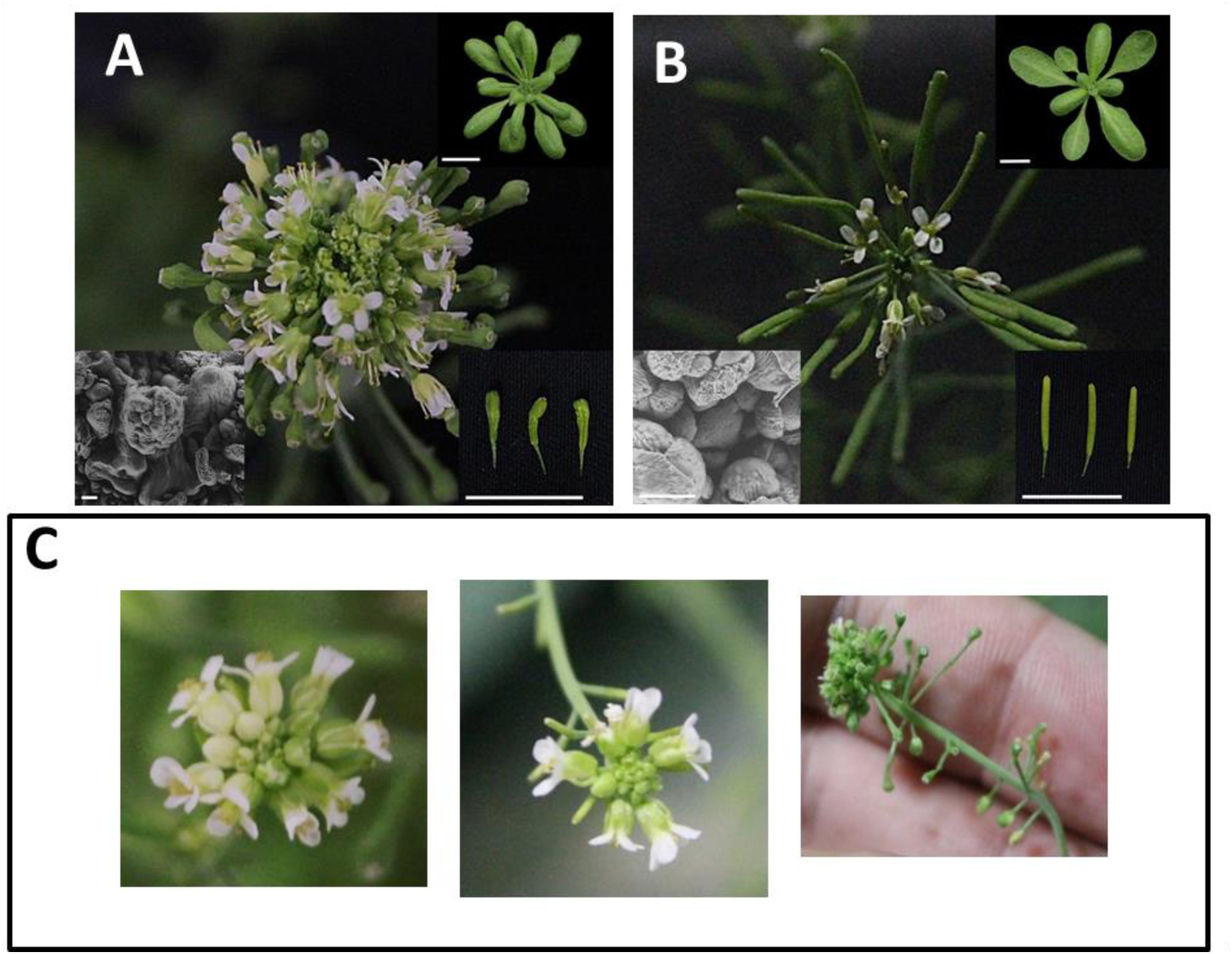
*S->C 9-3* ‘clv’ X *C9* F2 plants produced variable shoot phenotypes. (**A**) *S->C 9-3* ‘clv’ plants were crossed with (**B**) the *C9* progenitor line. The F1 plants appeared phenotypically ‘Wt.’ (**C**) The F2 progeny produced variable shoot morphologies that were not seen in either parent. Two other crosses likewise produced an F2 population with variable shoot phenotypes.

## Notes

### Competing Interest Statement

The authors have declared no competing interest.

